# A Co-essentiality Network of Cancer Driver Genes Better Prioritizes Anticancer Drugs

**DOI:** 10.1101/2024.12.20.629566

**Authors:** Kwanghwan Lee, Donghyo Kim, Inhae Kim, Juhee Lee, Doyeon Ha, Seongsu Lim, Eunjee Kim, Sin-Hyeog Im, Kunyoo Shin, Sanguk Kim

## Abstract

Diverse molecular networks have been extensively studied to discover therapeutic targets and repurpose approved drugs. However, it is necessary to select a suitable network since the performance of network medicine relies heavily on the completeness and characteristics of the selected network. Although a network using gene essentiality from cancer cells could be an effective platform for identifying anticancer targets, efforts to apply these networks in therapeutic applications have been limited. We constructed a phenotype-level network using the co-essentiality relationship between genes in CRISPR screens across 769 cancer cells to discover therapeutic targets for diverse cancer types. Leveraging cancer driver genes and network propagation on the networks, we found that the co-essentiality network better prioritized anticancer targets and biomarkers and predicted more precise drug responses in cancer cells than other molecular networks. The co-essentiality network outperformed conventional molecular networks in drug repurposing, which were validated in silico by clinical trial records. Notably, the co-essentiality network provided 37 repurposed drugs that the other networks have yet to cover, and we showcased three approved drugs repurposed for lung adenocarcinoma (Pioglitazone hydrochloride, Atovaquone, and Eflornithine). Our study provides a novel network for precision oncology to improve the identification of therapeutic targets in specific cancers.

## Introduction

Over the past few years, various anticancer drugs have been developed for diverse cancer types [1]. However, the overall clinical efficacy of approved drugs remains limited. Thus, identifying targetable alterations is urgently needed for the success of anticancer therapy. To achieve precision oncology, diverse tasks must be performed, such as identifying driver genes and discovering drug targets in specific cancer types.

Network-based approaches support precision oncology to identify robust anticancer targets or biomarkers linked to known disease genes [2,3], since genes related to disease phenotypes cooperate and cluster in the network [4]. We recently identified biomarkers of chemotherapy and immunotherapy by propagating the relatedness of therapeutic agents from drug targets to their neighbors in a protein-protein interaction (PPI) network [5,6]. Cheng et al. developed an *in silico* cancer drug repurposing framework using network modules derived from gene co-expression and PPI networks [7].

However, it remains to be seen which network is suitable for precision oncology, even though the choice of network is crucial to limit the performance of network-based approaches. It has been shown that network topology is a critical factor in improving the identification of disease genes. Huang et al. showed that the performance gap could be > 5,000× between the networks, from the highest to the lowest performance [8]. Buphamalai et al. also reported that each network is relevance to specific tasks [9]. Therefore, rather than relying on a one-size-fits-all network, we designed and evaluated networks that perform best on the tasks.

Co-essentiality networks, composed of genes with similar knockout essentiality profiles across various cancer cell lines, can be suitable networks to identify therapeutic targets in cancer. While several co-essentiality networks have been used for gene function prediction[10,11], the benefit of using co-essentiality networks in drug discovery still needs to be assessed. For example, it is reported that if two genes have similar essentiality profiles, they tend to have similar biological functions[10]. Co-essentiality networks have been applied to discover new functions of genes [11–13] and to infer genes into the same functional complexes[14] or metabolic pathways[15]. However, attempts to identify cancer drug targets using co-essentiality networks have been limited to discovering surrogate targets for several challenging target protein cases[12], despite the therapeutic opportunity that the essentiality phenotype might possess.

The potential of co-essentiality networks for drug repurposing remains particularly promising compared to conventional approaches. Most existing drug repurposing methods have relied on protein-protein interaction networks[7,16] or gene co-expression networks[17,18], which may have limitations in capturing functional relationships relevant to drug response. A co-essentiality network built from cancer cell line data may offer complementary insights by incorporating cellular fitness phenotypes. These phenotype-level relationships could provide additional perspective on drug effects compared to physical interactions or expression correlations. To our knowledge, this study represents the first comprehensive evaluation of co-essentiality networks for drug repurposing applications.

In this study, we aimed to assess diverse *in silico* frameworks using a co-essentiality network to identify novel anticancer drugs for specific cancer types and investigate the advantages of this network compared with conventional molecular networks. We find that the co-essentiality network was able to prioritize cancer-type-specific therapeutic targets and discover drug-repurposing candidates. The co-essentiality links formed highly clustered network modules of potential therapeutic targets. Moreover, the co-essentiality network predicted more precise drug responses in cancer cells than other molecular networks. We anticipate that the co-essentiality network will be a valuable resource for precision oncology and provide new therapeutic opportunities for cancer patients.

## Results

### Construction of the co-essentiality network for drug discovery in cancer

To construct a network for the identification of therapeutic targets and drug repurposing candidates of each cancer type, co-essentiality links were inferred from the correlation of the gene essentiality profiles of cancer cells (**Figure 1A**). A co-essentiality network was constructed with 18,119 genes and 8,105,180 co-essentiality links based on gene-level essentiality scores using a CRISPR screening dataset from the Cancer Dependency Map (DepMap) project [19]. Co-essentiality links represent the similarity of essentiality profile between two genes across cancer cells and are measured using the context likelihood of relatedness (CLR) algorithm [20]. To validate that the co-essentiality links from our method are relevant to biological function, we tested whether co-essentiality links are enriched in curated pathways. We found that the co-essentiality links indicate a strong functional relationship between the two genes (Figure 1B and S1), resulting in similar growth phenotypes across various cancer cells. For example, in KEGG pathways [21], gene pairs with greater link weights were likely within the same pathways (Figure 1B, blue points), and this tendency was higher than expected by chance (Figure 1B, gray points). Similarly, gene pairs with greater link weights likely reside within the same biological modules from five independent datasets: protein complexes (CORUM)[22], molecular pathways(REACTOME)[23], and Gene Ontology annotations[24] [GO: BP (biological process), MF (molecular function), and CC (cellular components)].

**Figure 1.**
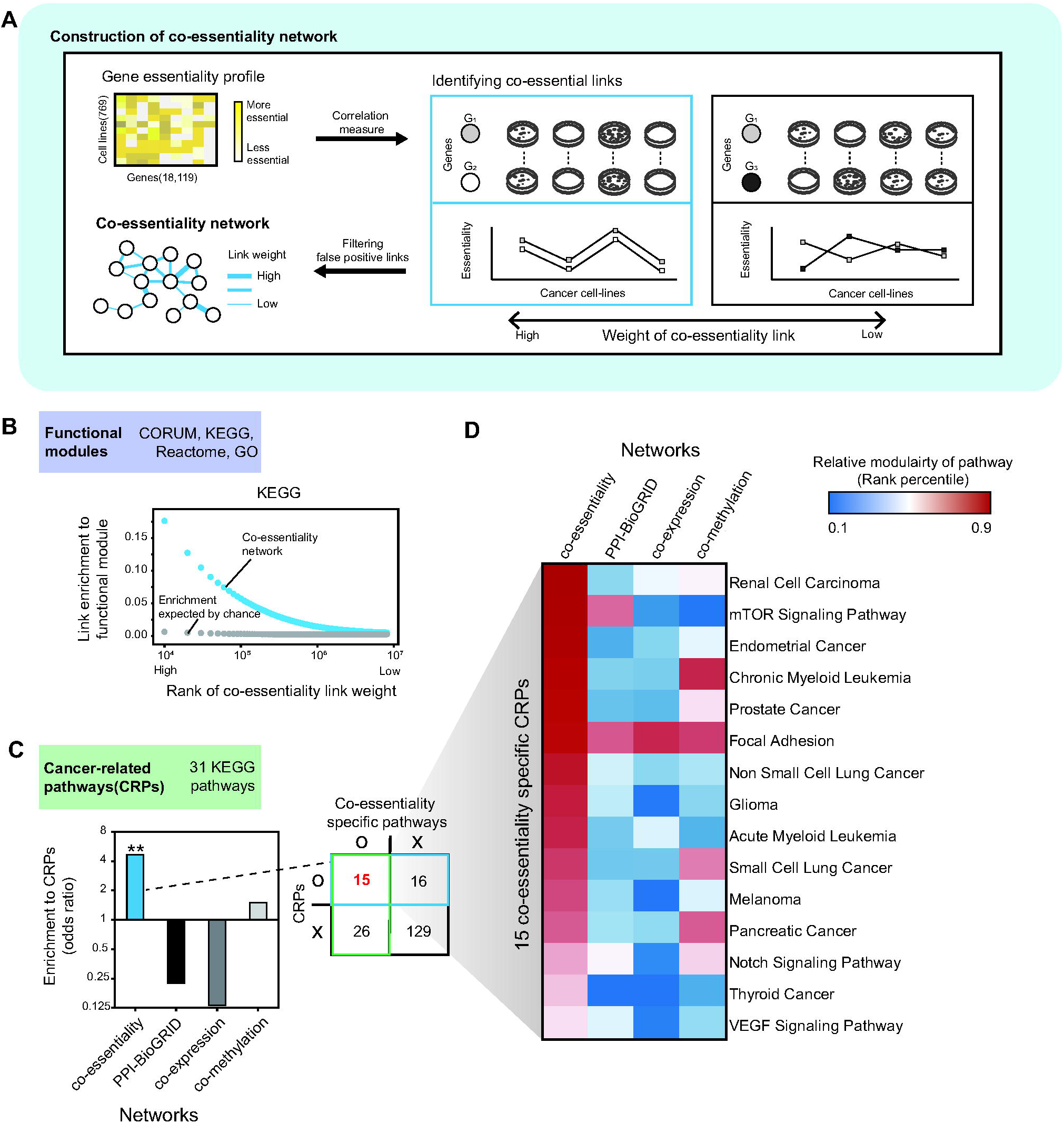
Construction of the co-essentiality network and validation for cancer association. **A.** Schematic illustration of constructing the co-essentiality network and its validation. CLR, context likelihood relatedness. **B.** Enrichment of the co-essentiality links to KEGG pathways. Co-essentiality links were ranked and binned (n = 10,000). Enrichment of the co-essentiality links in each bin is shown using blue dots. The expected enrichment of co-essentiality links in each bin is shown using grey dots. **C.** Enrichment of network links to 31 KEGG cancer-related pathways (CRPs) for four networks: the co-essentiality network, PPI network (BioGRID), co-expression network, and co-methylation network (Left). The contingency table of KEGG pathways with two criteria: inclusion in CRPs and inclusion in co-essentiality network-specific pathways. **D.** The relative modularity of 15 pathways overlapped between the 31 CRPs and the 41 co-essentiality-specific KEGG pathways.

### Effectiveness of the co-essentiality network in identifying cancer-related pathways

We found that the co-essentiality network depicts cancer-related pathways (CRPs) related to oncogenesis and hallmarks of cancer. To investigate the advantages of using the co-essentiality network, we compared the co-essentiality network with three other molecular network: a PPI network (BioGRID; 18,708 nodes; 434,527 links) [25] a co-expression network (19,120 nodes; 12,759,793 links), and a co-methylation network (16,333 nodes; 10,193,089 links). The co-essentiality network had a higher enrichment of CRPs (Figure 1c, left panel) with an odds ratio of 4.65, compared with the PPI network (odds ratio = 0.225), the co-expression network (odds ratio = 0.133), and the co-methylation network (odds ratio = 1.50). Co-essentiality links showed significant enrichment to CRPs (hypergeometric test p-value = 3.12×10^−3^), whereas other molecular links did not show significant enrichment to CRPs. Among 41 CRPs, the 15 CRPs had the highest relative modularity in the co-essentiality network (Co-essentiality specific pathways) and were greater than 6.83 CRPs, which was expected by chance (Figure 1C right, Table S1).

For example, in 1 of the 15 CRPs, the mTOR signaling pathway (KEGG: hsa04150), targeted by many anticancer drugs [26,27], the co-essentiality network showed higher modularity than the other networks (Figure 1D, rank percentile = 0.898, 0.697, 0.177, and 0.109 of relative modularity for the co-essentiality network, BioGRID, co-expression network, and co-methylation network, respectively). To determine whether co-essentiality links were likely to be found between gene pairs in CRPs, we measured the relative modularity of genes in each KEGG pathway and compared it with three other molecular networks (see Methods, Table S1). Since modularity is biased to degree of the nodes, we used degree-controlled modularity measure normalized by permutation test with degree matched random nodes. These results suggest that co-essentiality links can detect more oncogenic relationships compared with other molecular links, leading to improve identifying genes critical for cancer treatment.

### High modularity of cancer driver genes in the co-essentiality network

To verify the effectiveness of co-essentiality links for identifying therapeutic targets in cancer, we investigated the modularity of disease genes in the network, since the modularity of disease genes is indicative of the relevance underlying information to a particular disease phenotype [9]. We used cancer driver genes whose mutations cause cancer as disease genes. The co-essentiality links resulted in a denser module of cancer driver genes compared with other molecular links. For 17 out of 19 cancer types, the modularity (***m***) of driver genes was the highest in the co-essentiality network compared with the other three molecular networks (**Figure 2A**). Since modularity measures can be affected by the degree of nodes which is high in the co-essentiality network, we used more conservative measures, degree-controlled modularity(z-score) measure normalized by permutation test (see Methods). For instance, in the case of lung squamous cell carcinoma (LUSC), the co-essentiality network showed the highest modularity for driver genes (***m***_co-essentiality network_ = 15.61) among all molecular networks (***m*** _PPI-BioGRID =_ 3.38, ***m*** _co-expression network_ = 3.33, and ***m*** _co-methylation network_ = 1.99) (Figure 2B-E, and S2). For example, comparing the driver genes of LUSC (Figure 2B) among the networks (Figure 2C-E), the subnetwork of driver genes in the co-essentiality network was more densely connected. *FAT1* and *FGFR2* connect to the subnetwork of LUSC driver genes in the co-essentiality network but are not connected to other molecular networks. *FAT1* has co-essentiality interactions with *RASA1*, *CUL3*, and *ARHGAP35*. *FGFR2* has co-essentiality interaction with *FAT1*. Indeed, *FAT1* is a biomarker of immune checkpoint blockade for lung cancer [28], and inhibition of *FGFR2* is also effective for the treatment of lung cancer [29]. The 17 cancer types with the highest modularity in the co-essentiality network were selected for further analyses of precision oncology tasks.

**Figure 2.**
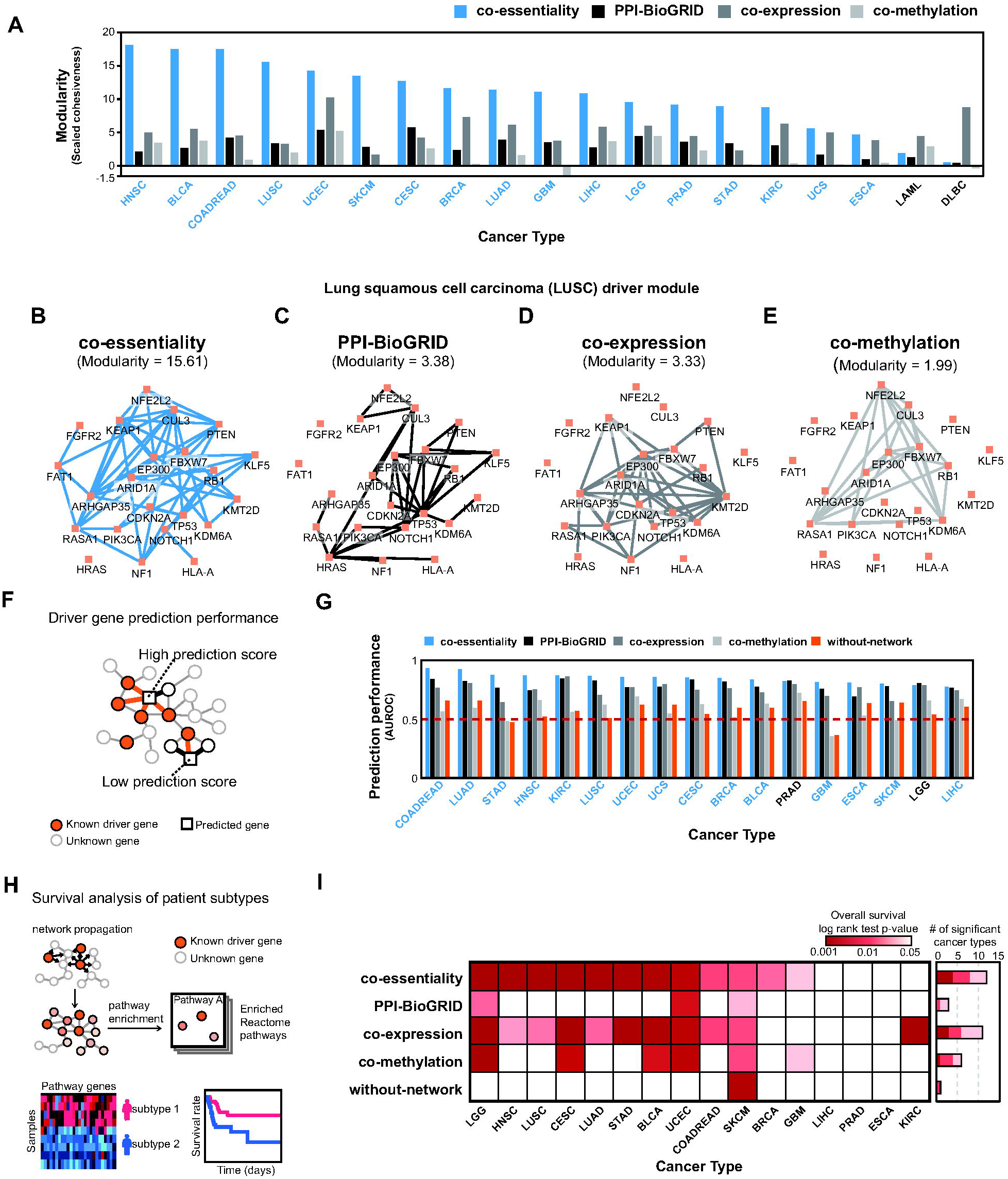
High modularity of cancer driver genes in the co-essentiality network is the key to driver gene identification and patient stratification. **A.** Modularity calculated from a subnetwork of driver genes across 19 TCGA cancer types in four networks: co-essentiality, PPI-BioGRID, co-expression, and co-methylation. For blue-colored cancer types, the co-essentiality network showed the highest modularity among the four networks. **B-E.** Illustrations of the subnetwork of lung squamous cell carcinoma (LUSC) driver genes in each network. **F.** A schematic illustration of driver gene identification using the co-essentiality network. **G.** Driver gene identification performance in the four networks(co-essentiality; PPI-BioGRID; co-expression; co-methylation), and the without-network control. For blue-colored cancer types, the co-essentiality network showed the best performance among the four networks. **H.** A schematic illustration of TCGA patient stratification using the co-essentiality network. **I.** Results for patient stratification using the four networks, and the without-network control. A heatmap showing the significance of survival difference between two patient groups stratified by each network tested using the log-rank test.

To investigate the robustness of the results, we leveraged a different driver gene resource, Cancer Gene Census (CGC) [30], the modularity (***m***) of driver genes was the highest in the co-essentiality network in 12 out of the 19 cancer types (Figure S3; see Methods). Additionally, we employed a distinct modularity metric, the clustering coefficient, which focuses more on link connectivity. For 16 out of the 19 cancer types, the clustering coefficient of driver genes in the co-essentiality network was higher than that in other molecular networks (Figure S4). We also explored how the modularity of driver genes is influenced by the total number of cell lines used to construct the co-essentiality network. With the inclusion of more cancer cell lines, the co-essentiality networks became more refined, revealing additional disease modules that could serve as potential therapeutic targets (Figure S5).

We further investigated whether the high modularity of co-essentiality links could be utilized to identify the driver genes of each cancer type via connection with known driver genes based on guilt-by-association (Figure 2F). We found that co-essentiality links are more suitable than other molecular links for identifying cancer driver genes. The co-essentiality network outperformed the three other molecular networks in discovering driver genes using guilt-by-association in 15 of the 17 cancer types (Figure 2G). For example, in the case of head and neck cancer, co-essentiality links improved driver gene discovery by 17%, 16%, and 33% than other networks, respectively. Similarly, the co-essentiality network with guilt-by-association showed higher performance in 13 of the 17 cancer types compared with seven other PPI networks (Figure S6): BioPlex[31], GPSnet [7], HURI [32], InBioMap [33], iRefIndex [34], Pathway Commons [35], and STRING [36]. Furthermore, our co-essentiality network demonstrated superior performance in identifying driver genes compared to previously published co-essentiality networks (Wainberg_etal [11], Amici_etal [12], and Gheorghe_etal_Ceres, and Gheorghe_etal_BF [37]) and a genetic interaction network based on synthetic-lethal relationships [34] (cSLnet) in 14 out of 17 cancer types (Figure S7). The co-essentiality network also surpassed a ‘without-network’ control method that relied solely on the gene essentiality of cell lines from a specific cancer type, devoid of network support (see Figure 2G, orange bars).

One might wonder if the performance of the co-essentiality network is driven by the frequency of tumor types used to construct it. We found that the ability of the co-essentiality network to identify driver genes had no significant correlation with the number of cell lines per tumor type in the DepMap (Spearman correlation coefficient R = 0.167, p-value = 0.522, Figure S8). Additionally, we evaluated if the characteristic of the co-essentiality network was merely influenced by a high proportion of essential genes within it. Contrary to what might be expected, only 11.4% of genes in the co-essentiality network are essential, placing it 8th among the 13 networks (Table S2).

Using two additional reported methods for driver gene identification, we found that co-essentiality links resulted in the best performance improvements and had a robust benefit for the identification of cancer driver genes. The performance for driver genes identification was measured using network propagation methods such as Hotnet2 [38] (Figure S9A) and uKIN [39] (Figure S9B). In both methods, co-essentiality links identified driver genes better than any other molecular links. For 6 out of 11 cancer types used in the Hotnet2 study [38], the co-essentiality networks with the Hotnet2 algorithm outperformed 15 other networks in identifying driver genes (Figure S9C). Compared to the co-expression network, which showed the second-highest performance, the co-essentiality network achieved significantly higher F1 scores with no significant correlation (Wilcoxon signed-rank test p-value = 2.93 × 10-3; Pearson correlation coefficient[PCC] p-value = 0.0613; Figure S9E and F). Similarly, when using the uKIN algorithm, the co-essentiality network ranked among the top 3 performers in 15 of the 24 cancer types examined in the uKIN study [39] (Figure S9D). Although the Amici_etal network ranked in the top 3 for 19 of the 24 cancer types, there was no significant difference between their actual AUROC values (Wilcoxon signed-rank test p-value = 0.169; Figure S9G). Furthermore, the AUROC rankings of these two networks showed inverse correlation (PCC = −0.452, p-value = 0.0266), indicating that each co-essentiality network with the uKIN algorithm exhibits distinct strengths across different cancer types. Taken together, our results that the co-essentiality network captures more relationships between cancer driver genes suggest that co-essentiality links are more relevant to cancer phenotypes than other molecular links.

### Driver modules of the co-essentiality network can differentiate patient survival

To examine the clinical relevance of the modularity of driver genes in the co-essentiality network, we evaluated whether the network modules derived from these driver genes improve our understanding of cancer prognosis in terms of patient survival (Figure 2H). Among the 17 TCGA cancer types, 16 with enough patients’ survival data were tested. We found that driver modules from the co-essentiality network were more informative for stratifying patients along with overall survival than other molecular networks (Figure 2I and S10). In each network, driver modules representing biological pathways in significant network proximity to the driver genes were tested to discern patient survival. Specifically, patients were divided into two groups based on the overall expression of the driver module, and the significance of the survival difference between the two groups was measured using the log-rank test.

The co-essentiality links provided the best driver modules that stratified patients into groups with different survival rates in 12 of 16 cancer types. In contrast, the driver modules from other molecular networks failed to stratify patients in 13, 6, and 10 cancer types for PPI (BioGRID), co-expression, and co-methylation networks, respectively (Figure 2I and S10-13). For example, in LUSC, the pathway of Signaling by Non-Receptor Tyrosine Kinases (R-HSA-9006927), which is significantly proximal to driver genes in the co-essentiality network (normalized enrichment score [NES] = 4.13, false discovery rate [FDR] = 9.99×10^−4^; Figure S15A), can stratify patients with its down-regulation, exhibiting longer overall survival than the others (p-value = 3.75×10^−4^, log-rank test; Figure S15B and S15C). In contrast, co-expression and co-methylation links lined up different driver modules, Ovarian tumor domain proteases (R-HSA-5689896) and Signaling by FGFR2 in disease (R-HSA-5655253), respectively, and failed to differentiate the overall survival of patients (p-value = 0.0255, co-expression; p-value = 0.314, co-methylation, figs. S12 and S13). PPIs provided no driver module because no biological pathway was significantly proximal to driver genes (Figure S11).

We also found that driver modules based on co-essentiality links were more informative for patient survival than the driver genes themselves. The control method using driver genes alone, without module identification using the co-essentiality network, was unable to differentiate survival outcomes in 15 out of 16 cancer types (bottom line of Figure 2I and S14). In LUSC, for example, the patient groups stratified by expression levels of the driver module using co-essentiality links, Non-Receptor Tyrosine Kinases pathway, showed a significant difference in the overall survival (Figure S15C, p-value = 3.75×10^−4^); in contrast, the patient groups distinguished by the expression level of driver genes themselves showed no significant difference in overall survival (Figure S15D, [LUSC]; p-value = 0.231).

### Co-essentiality network can identify potential drug targets or biomarkers for specific cancer types

To explore the therapeutic use of the co-essentiality network, we examined the potential of co-essentiality links for drug repurposing tasks in cancer. Using network propagation, we examined genes closely located to driver genes in the co-essentiality network. These genes may serve as an anticancer drug’s targets or biomarkers (i.e. drug-associated genes [DAGs]). We utilized them for identifying new indications of approved drugs (**Figure 3A**). We verified the performance of the co-essentiality network in identifying therapeutic target genes by three ways: (a) prioritization of FDA approved drug-associated genes(DAGs) [40] (Table S3); (b) prediction of cytotoxicity of drugs on cancer cells (IC_50_); (c) prediction of reversal gene expression (RGE) effect of drugs on cancer cells [41]. Finally, we validated 145 repurposing candidates, predicted through co-essentiality links, using cancer-related clinical trial records.

**Figure 3.**
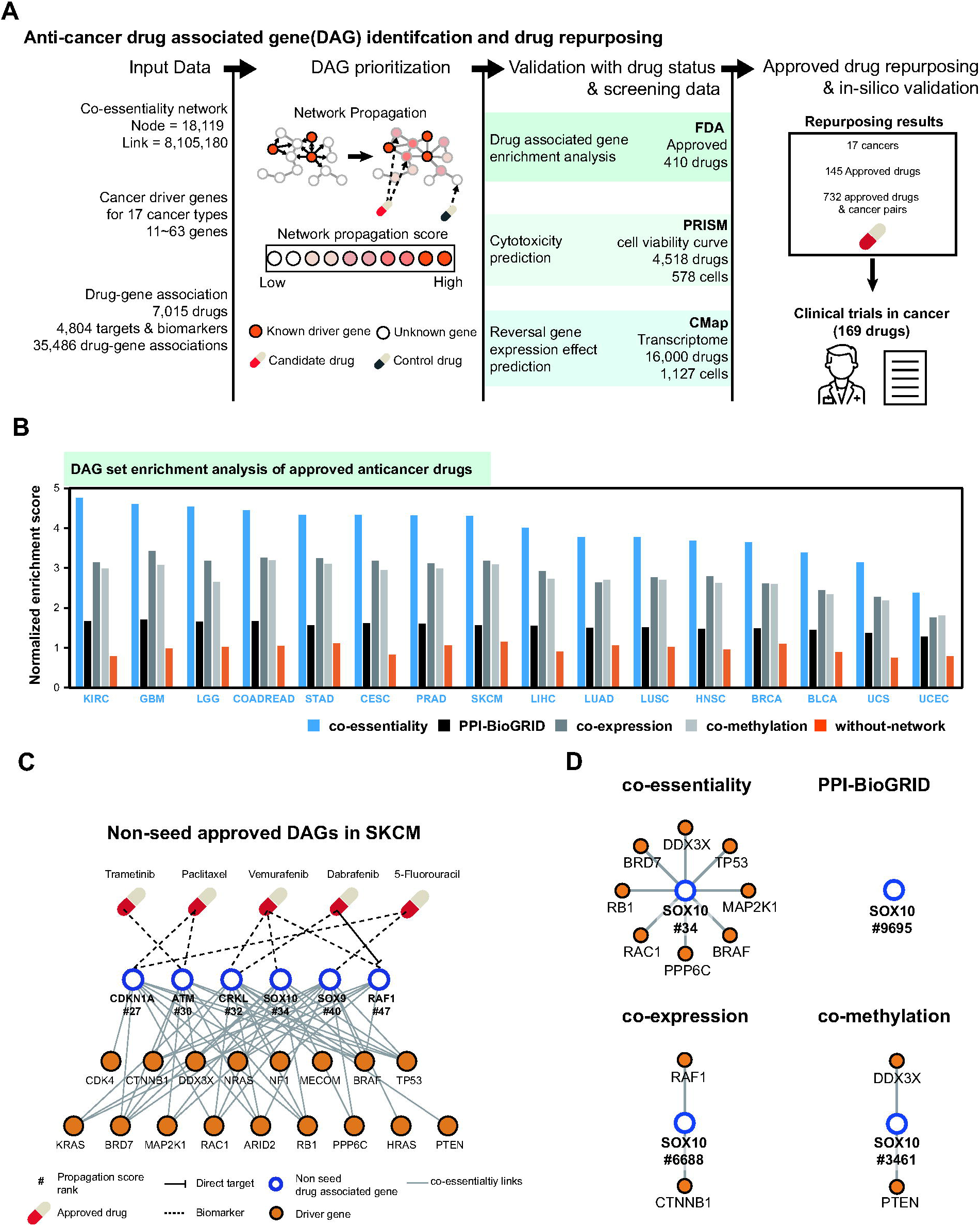
Prioritization of anticancer drug-associated genes in 16 cancer types using the co-essentiality network. **A.** A schematic view of drug-associated gene(DAG) prioritization using the co-essentiality network and the validation of prioritized DAGs. **B.** Normalized enrichment score (NES) from Gene Set Enrichment Analysis(GSEA) of the four networks(co-essentiality, PPI-BioGRID, co-expression, and co-methylation), and the without-network control. For blue-colored cancer types, the co-essentiality network showed the highest NES among the four networks. **C.** Subnetwork of six approved DAGs, *CDKN1A*, *ATM*, *CRKL*, *SOX10*, *SOX9*, and *RAF1*) in SKCM and driver genes in their first neighbors in the co-essentiality network. Only drug-gene associations of these six genes are shown in this figure. **D.** First neighbors of *SOX10* in the four networks.

Network propagation with co-essentiality links better prioritized DAGs of FDA-approved anticancer drugs compared with 15 other networks including other co-essentiality networks(Figure 3B, S16A, and S17A). We investigated 16 out of the 17 TCGA cancer types, excluding ESCA, which lacks available FDA-approved DAGs. In all 16 of these cancer types, the propagation values, determined using co-essentiality links, demonstrated the highest positive enrichment for the FDA-approved DAGs. The average normalized enrichment score (NES) of co-essentiality links was 4.14, 39% higher than that of co-expression links that had the second-highest NES (Figure 3B). We also performed separate analyses for drug targets and biomarkers. For drug targets, analysis was limited to 9 out of 17 TCGA cancer types due to a lack of approved target information for the remaining cancer types. The co-essentiality network outperformed other networks in prioritizing targets for 5 of these 9 cancer types (Figure S16B, S17B, and S18A). For biomarker prioritization, the co-essentiality network demonstrated superior performance across all 15 analyzable cancer types (Figure S16C, S17C, and S18B). Having greater NES to propagation values of co-essentiality links suggests that the co-essentiality network prioritizes DAGs of anticancer drugs better. For example, in skin cutaneous melanoma (SKCM), the co-essentiality network had an NES value of 4.18, which was higher than those of PPI-BioGRID, co-expression network, and co-methylation network (1.59, 3.25, and 3.08 respectively). Among the top 50 genes with the highest propagation value from the co-essentiality network, 20 were identified as approved drug-associated genes which were higher than the number of PPI-BioGRID, co-expression network, and co-methylation network (18, 14, and 14 genes in the top 50 genes with the highest propagation value, respectively). Among these 20 genes, 3 (*BRAF*, *MAP2K1*, and *RAF1*) were both targets and biomarkers and 17 (*TP53, RAC1, CTNNB1, KRAS, RB1, PTEN, NF1, NRAS, HRAS, KIT, GNA11, CDKN2A, CDKN1A, ATM, CRKL, SOX10,* and *SOX9*) were only biomarkers of approved drugs (Table S4).

Furthermore, we found that the co-essentiality network outperformed the control method which has no the assistance of network propagation (Figure 3B, orange bars). Specifically, the co-essentiality network identified six genes (RAF1, CDKN1A, ATM, CRKL, SOX10, and SOX9) due to their multiple connections with SKCM driver genes employed as initial inputs for propagation, despite not being designated as seed genes themselves (Figure 3C). RAF proto-oncogene serine/threonine-protein kinase (*RAF1*) is one of the targets of Dabrafenib, a FDA-approved *BRAF* inhibitor[42,43]. In the co-essentiality network, *RAF1* showed a high propagation value (1.37 × 10^−4^ and ranked 47th) because *RAF1* is linked to five known SKCM driver genes: *CTNNB1, NRAS, BRAF, NF1,* and *KRAS*. Similarly, *ATM*, biomarker of Paclitaxel[44,45] and Trametinib[46], is linked to eight known SKCM driver genes: *BRD7*, *MAP2K1*, *CTNNB1*, *RB1*, *ARID2*, *BRAF*, *DDX3X*, and *TP53*. These results represent the advantages of utilizing the co-essentiality network approach over the gene-centric approaches without network.

We observed that co-essentiality links could identify the approved drug-associated genes that were not captured by other molecular links. For example, in SKCM, a biomarker of Vemurafenib[47], *SOX10*, ranked 34^th^ in the co-essentiality network, but 9,695^th^ in the PPI network, 6,688^th^ in the co-expression links, and 3,461^st^ in the co-methylation network (Figure 3D). The co-essentiality network can identify more interactions between *SOX10* and known driver genes of SKCM that are not captured in other molecular links. *SOX10* has co-essentiality links with seven driver genes of SKCM: *TP53, MAP2K1, BRAF, PPP6C, RAC1, RB1*, and *BRD7*, which are not connected to other molecular networks. This suggested that co-essentiality links could help to facilitate the identification of new drug-associated genes which are not covered by other molecular links.

### Co-essentiality network finds candidates for drug repurposing

Having confirmed the capability of co-essentiality links to find DAGs of approved drugs, we further investigated whether they could facilitate network-based prediction of drug responses assessed in large-scale pharmacogenomic screenings. For each drug, we assigned a therapeutic candidate (TC) score aggregated from the propagation value of its DAGs by taking their root mean square (RMS). Therefore, we considered the average effect of DAGs on specific cancer types to eliminate bias to the number of DAGs.

We found that the TC score calculated from co-essentiality links predicted the cytotoxicity of drugs for cancer cells better than the other molecular links. For each cancer type, drugs with a high TC score from the co-essentiality links were highly cytotoxic to cancer cell lines from that cancer type. Co-essentiality links showed the highest correlation between TC score and the IC_50_ in 13 out of 15 cancer types which have drug screen data (**Figure 4A**), and the average increase in the Spearman correlation coefficient (Spearman R) was 9.7%, 8.4%, and 43% for the PPI, co-expression, and co-methylation links, respectively. For example, TAK-733, an MEK1/2 inhibitor under clinical trials for advanced non-hematologic malignancies and advanced metastatic melanoma [48,49], was predicted to have an antitumor effect on colorectal cancer (COADREAD; Figure 4B; TC score = 5.39×10^−3^, rank percentile = 98.57%), and indeed showed high cytotoxicity in the cancer cell lines of COADREAD (median IC_50_ = 1.03 × 10^−3^ µmol). In the co-essentiality network, TAK-733 had seven DAGs, *MAP2K1, MAP2K2*, *BRAF, KRAS, NRAS, PIK3CA,* and *GNA11* (Figure 4C, red circles), which are interconnected by nine driver genes of colorectal cancer, namely *CTNNB1, SOX9, TP53, APC, TCF7L2, TGF1, FBXW7, PTEN,* and *ARID1A* (Figure 4C, orange filled). Similarly, the co-essentiality network showed a higher correlation between the TC score and IC_50_ in 12 out of 15 cancer types than seven other PPI networks (Figure S19). Similarly, the co-essentiality network showed a higher correlation than other co-essentiality networks and a genetic interaction network in 8 out of 15 cancer types (Figure S20).

**Figure 4.**
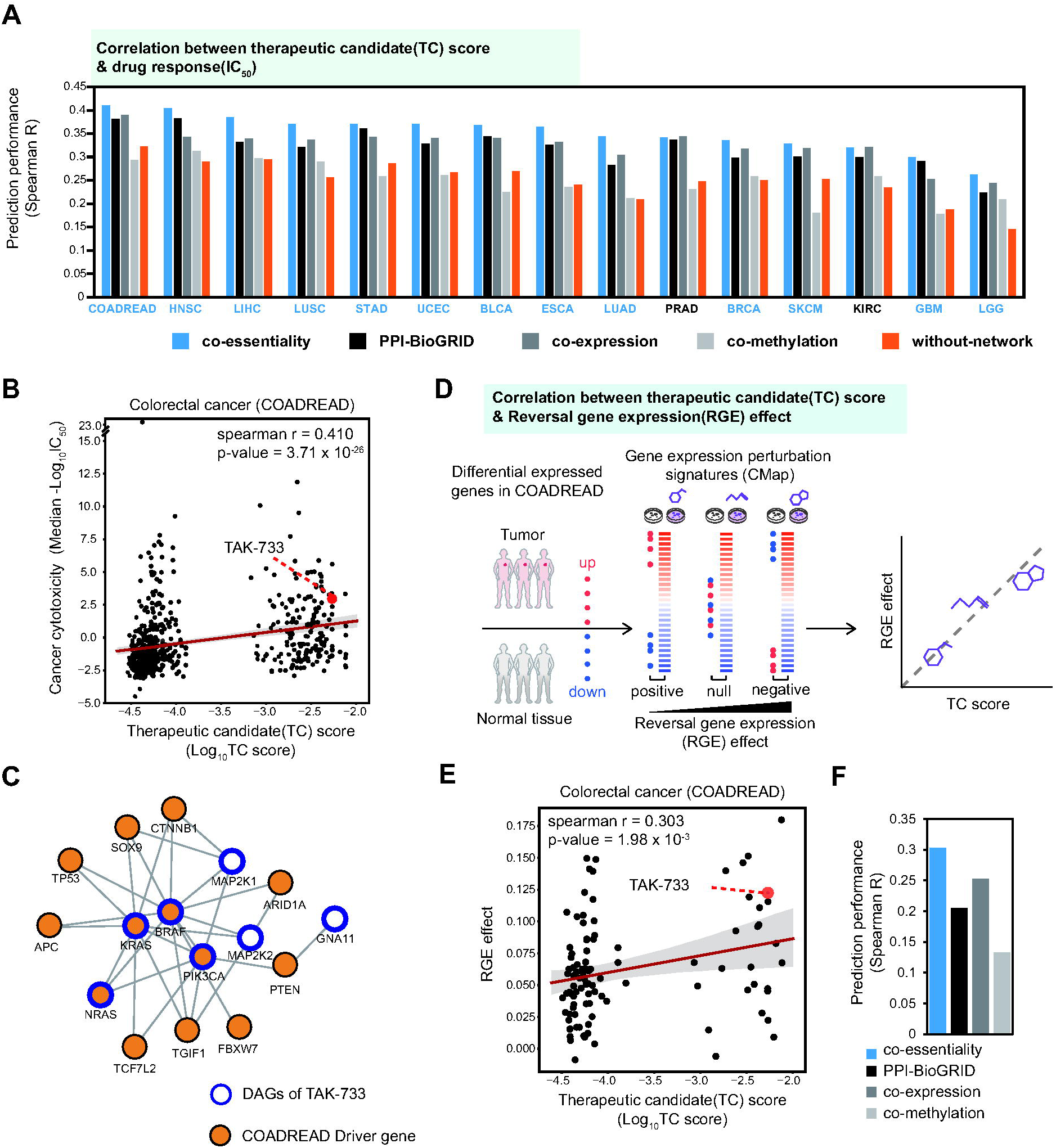
Predicting drug response in cancer cells of 15 cancer types using therapeutic candidate (TC) score of the co-essentiality network. **A.** Prediction performance of drug response measured using the Spearman correlation coefficient between TC score and median -log_10_(IC_50_) for the four networks(co-essentiality, PPI-BioGRID, co-expression, and co-methylation), and the with-out network control. For blue-colored cancer types, the co-essentiality network showed the best performance among the four networks. **B.** Scatter plot of drug response and TC score of colorectal cancer in the co-essentiality network and data point of candidate drug: TAK-733 (red dot). **C.** A subnetwork of seven DAGs of TAK-733 and COADREAD driver genes which are connected to them in the co-essentiality network. **D.** Schematic diagram of drug’s reversal gene expression (RGE) effect on COADREAD. **E.** Scatter plot of drug’s RGE effect and TC score of COADREAD in the co-essentiality network. The red data point is a candidate drug, TAK-733. **F.** Performance of the four networks for predicting drug RGE effect in COADREAD.

One might ask whether driver genes alone, without the assistance of network propagation, can predict drug responses. We found that the control method using driver genes alone was not as predictive as the TC score of the co-essentiality network (Figure 4A, orange bars). The TC score showed a greater correlation with IC_50_ for all 15 cancer types than driver genes, and the average increase in the Spearman correlation coefficient was 42%. This is because driver genes cover only 4.4% of drug targets, whereas using network propagation, all the genes in networks were assigned propagation scores.

Additionally, we assessed whether incorporating biomarker information enhances the accuracy of drug response predictions. The TC score using DAGs including both target and biomarker information showed a greater correlation with IC_50_ across 15 cancer types than using only target information, and the average increase in the Spearman R was 32.1% (Figure S21). This result indicates that leveraging both drug targets and biomarkers improves predicting the anticancer effects of drugs, compared to relying solely on direct target data.

Focusing on COADREAD, as it showed the greatest correlation between TC score and IC_50_, we further discovered that drugs with a high TC score could alter gene expression to revert to that in cancer conditions (Figure 4D). We used the degree of reversal effect of the drugs on cancer-associated gene expression to confirm the validity of the TC score, since a drug capable of ameliorating the perturbed molecular state induced by a disease exhibits potential for therapeutic efficacy. We measured the reversal gene expression (RGE) effect [50], which takes the dot product of ranked differential expression between cancer [51] and drug-treatment conditions[52] (Figure 4D), and investigated its association with the TC score. We observed that the TC score of the co-essentiality network was positively correlated with the RGE effect (Figure 4E; Spearman R = 0.303, p-value = 1.98×10^−3^). For example, TAK-733, which exhibited a high TC score and -log_10_(IC_50_) in Figure 4B, showed a strong reversal effect on the expression pattern (RGE effect = 0.123, rank percentile = 91.18 %). Additionally, the TC score from the co-essentiality network exhibited a greater correlation with the RGE effect than other molecular networks (Figure 4F). Overall, these results indicate that the co-essentiality network is a more effective than the molecular networks based on PPI, co-expression, and co-methylation for finding driver-associated therapeutic targets, which holds potential for drug repurposing.

### In silico drug repurposing of approved drugs using the co-essentiality network

To identify the novel therapeutic potential of the approved drugs for other purposes, we conducted *in silico* drug repurposing using the TC score from the co-essentiality network. Among a total of 1,213 approved non-cancer drugs having drug-gene associations from the PanDrugs database [53], 145 were predicted to have significant anticancer activities in at least one of the 17 cancer types by the co-essentiality network (**Figure 5A**; adjusted P < 0.05; Table S5). To find repurposing candidates, we measured the significance of the TC score of drugs from permutation test in each cancer type (See methods). For example, Pioglitazone hydrochloride has the TC score = 1.3 × 10^−3^ in LUAD, and the significance of the TC score calculated from the permutation test of random DAGs is adjusted P = 3.79 × 10^−3^ (Figure S22).

**Figure 5.**
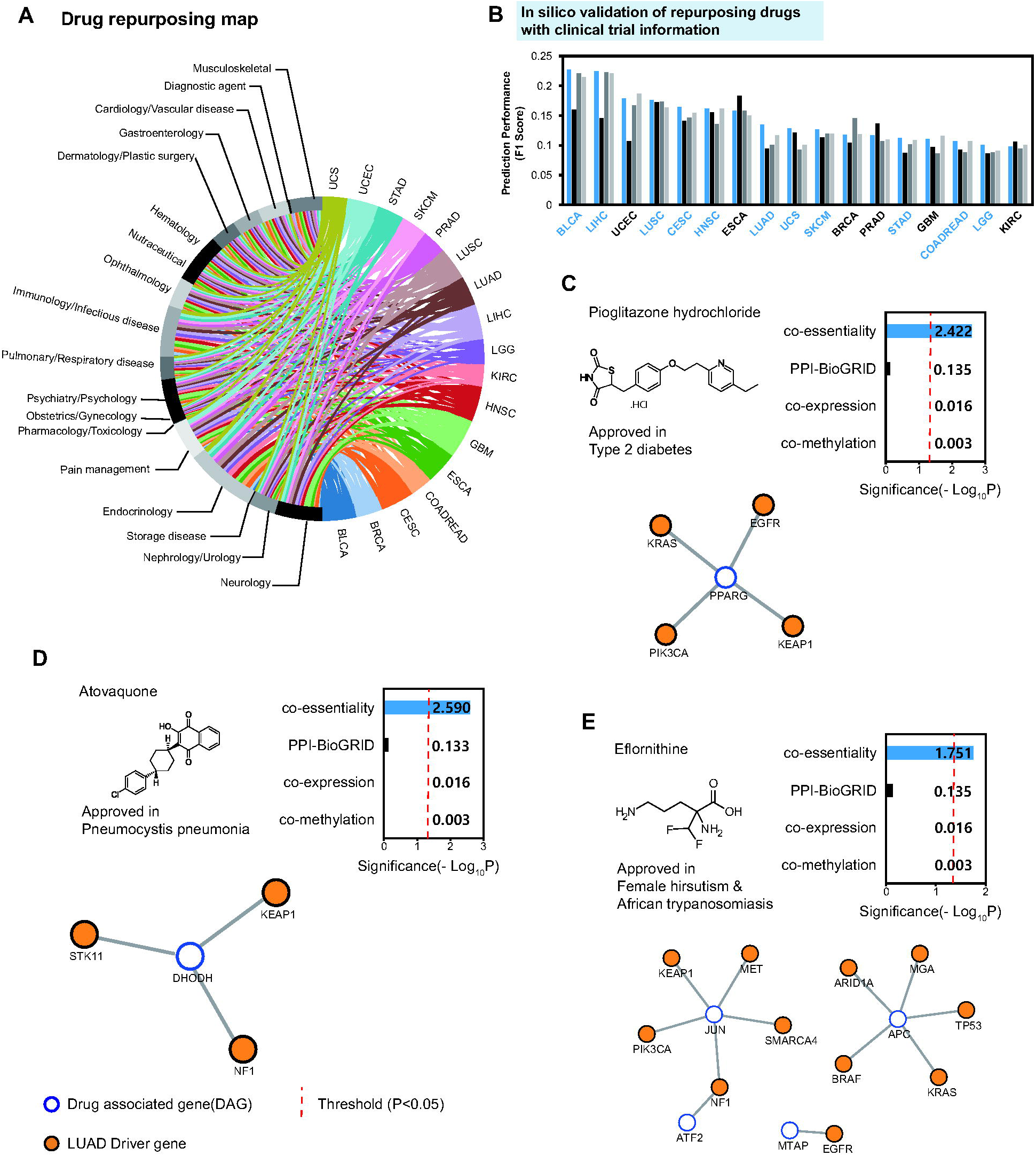
In silico drug repurposing using co-essentiality network and case studies in LUAD. **A.** A chord plot illustrating a global view of potential anticancer indications for 145 approved drugs across 17 cancer types. B. Validation of the drug’s new indication using clinical trial records. The prediction performance of drug indication is measured with the F1 score. For blue-colored cancer types, the co-essentiality network showed the best performance among four networks: co-essentiality, PPI-BioGRID, co-expression, and co-methylation. C-E. Drug repurposing candidates in LUAD, which is predicted by the co-essentiality network but not by other networks. The bar graphs showed the significance of the TC score and the red dot lines showed the threshold of drug repurposing (adjusted P=0.05). In the below, a subnetwork of drug-associated genes (DAGs) and LUAD driver genes are connected to them in the co-essentiality network.

To further assess the effectiveness of *in silico* drug repurposing utilizing the co-essentiality network, we explored whether the repurposing candidates had clinical trial records related to cancer. For 11 out of the 17 cancer types analyzed, the repurposing results from the co-essentiality network demonstrated enhanced performance in predicting drugs having clinical trials for cancer, compared to that from PPI-BioGRID, co-expression, and co-methylation networks (Figure 5B). Similarly, the co-essentiality network showed a higher performance in 6 cancer types than seven other PPI networks (Figure S23). Also, the co-essentiality network showed a higher performance than four other co-essentiality networks and a genetic interaction network across seven cancer types (Figure S24).

Furthermore, we demonstrated that our co-essentiality network-based repurposing method outperformed an existing network-based approach presented in Cheng et al.[16], which has been widely applied across multiple diseases[54–56]. The Cheng et al. method relies on network proximity between disease genes and drug targets to generate drug repurposing prediction scores. Across all 17 cancer types analyzed, our co-essentiality network showed superior performance in predicting drugs with cancer-related clinical trials compared to the Cheng et al. method, regardless of which of three proximity thresholds(z) was applied (Figure S25).

Specifically, we identified 37 non-cancer drugs that were given new indications not addressed by other networks. In the case of LUAD, we identified seven repurposing candidates solely predicted by the co-essentiality network, distinct from other networks, and interestingly, three of these drugs were associated with documented clinical trials in cancer (Table S6). For example, Pioglitazone hydrochloride, an antihyperglycemic, indicated to manage type 2 diabetes mellitus, was exclusively identified by the co-essentiality network (adjusted P = 3.79 × 10^−3^, Figure 5C). Pioglitazone is a selective agonist at peroxisome proliferator-activated receptor-gamma (*PPARG*) having co-essentiality links with four LUAD driver genes(*KRAS*, *EGFR*, *PIK3CA*, and *KEAP1*). This prediction is supported by both preclinical studies which investigated the effect of pioglitazone in the non-small cell lung cancer (NSCLC) cell lines, patients derived cancer cells[57], and NSCLC mouse models[58] Also, five clinical trials of Pioglitazone have been reported in lung cancer (NCT00780234, NCT05919147, NCT00923949, NCT02852083, and NCT01342770), and NCT05919147 is still in recruiting status now.

Another repurposing candidate, Atovaquone, a hydroxynaphthoquinone, is approved in Pneumocystis pneumonia and was predicted to have the significant antitumor effect on LUAD (P = 2.57 × 10^−3^, Figure 5D). Atovaquone inhibits a mitochondrial protein, Dihydroorotate Dehydrogenase(*DHODH*) which has co-essentiality links with three LUAD driver genes(*STK11*, *KEAP1*, and *NF1*), resulting in inhibition of the electron transport chain and a loss of mitochondrial function. Several studies have described the anticancer effect of atovaquone[59–62], and there are two clinical trials of atovaquone in non-small-cell lung cancer(NCT04648033 and NCT02628080).

The co-essentiality network found that Eflornithine, ornithine decarboxylase(*ODC1*) inhibitor, which was primarily used to treat female hirsutism and African trypanosomiasis was associated with LUAD (P = 0.0177, Figure 5E). Although in the co-essentiality network, *ODC1* has no connection with driver genes of LUAD, four biomarkers(*JUN*, *APC*, *ATF2*, and *MTAP*) have links with 11 LUAD driver genes. It is also supported by previous literature[63,64] and clinical trials(NCT05500508). These results suggest repurposing candidates using network propagation with the co-essentiality links can provide new therapeutic options.

## Discussion

In this study, we showed that therapeutic targets of cancer could be identified using co-essentiality links (Figure 3). We applied a co-essentiality network to systematically prioritize anticancer targets and biomarkers in various cancer types, and demonstrated that it performed better than other molecular networks, including PPI, co-expression, and co-methylation networks (Figure 3A). We constructed a co-essentiality network using phenotype information, where links were inferred from the correlation of the gene essentiality profile of cancer cells (Figure 1). Since gene essentiality is measured by proliferation rates of cancer cells [65], a crucial indicator of anticancer therapy, gene essentiality profiles have been applied to identify promising targets of anticancer therapies [19,66,67]. Gene essentiality was also utilized to form a network. For example, co-essentiality networks have been built to identify a group of genes with the same functional complexes [14] or metabolic pathways [15] and new functions of genes [10–13,68]. However, for therapeutic purposes, co-essentiality networks have not been applied to identify the target genes of anticancer therapies. Here, we demonstrated that links of the co-essentiality network could capture functional pathways related to the proliferation of cancer cells (Figure 1C and 1D). Our results suggested that co-essentiality links can prioritize DAGs for cancer treatment.

Since the network approaches are dependent on the topology of the networks, finding a suitable network is crucial for developing network medicine [8,9]. Here, we observed that the co-essentiality network showed an improvement in driver gene identification when compared with diverse molecular networks (Figure 2G). Using well-known network propagation approaches, such as Hotnet2 [38] and uKIN [39], our co-essentiality network showed improved prediction performance in identifying driver genes compared with other molecular layered networks (Figure S9). Specifically, the co-essentiality network showed higher modularity between driver genes than any other molecular network (Figure 2A, S3, and S4). This topological characteristic of the co-essentiality network contributes to the high performance of the driver gene identification. In fact, the modularity of disease genes in specific networks may be the key to identifying disease genes in various diseases [9]. Thus, for precision oncology, the choice of a network with highly clustered cancer driver genes has the potential to improve task performance.

It is essential to note that for each different purpose one might need to employ a customized approach to obtain an optimal network for that purpose. For the repurposing of drugs, we demonstrated that the co-essentiality network developed here performed better than other networks, including other co-essentiality networks (Figure S17, S20, and S24). Since the repurposing process relied on network propagation, our co-essentiality network would be taking advantage of the greater number of direct links than other networks, having much smaller loss of information during the propagation process. This observation aligns with earlier findings that the performance of disease gene identification was positively correlated with the size of a network [8]. Specifically, those direct links likely embody the phenotypic associations recorded from CRISPR-CAS9 screening into topological paths between the target and seed nodes[69]. By contrast, when aiming to discover novel biochemical interactions, such as metabolic pathways [15] or protein complexes [14], it would be more relevant to obtain smaller co-essentiality networks in which links have greater chance to represent such molecular interactions with less false positives. Given a specific purpose, it remains a crucial scientific challenge to identify an apt approach to building biological networks with varying link properties and topological features of the systems[9,70–72].

For drug repurposing tasks, we found that drug-gene association that includes both targets and biomarkers provided more benefit to predict the drug’s therapeutic potential. ‘Biomarker’ is a gene whose genetic status is linked to treatment response (based on pre-clinical or clinical evidence), but the protein product itself is not the pharmacological target. However, numerous studies have highlighted the potential for biomarkers to become druggable, and the ability to guide right drug-disease associations [53,73–77]. We discovered that utilizing drug-gene associations, which include both targets and biomarkers, improves the precision in predicting the anticancer effects of drugs over just using direct target data (Figure S20). Furthermore, our study identified Eflornithine as a potential candidate for repurposing, due to its four biomarkers (*JUN, APC, ATF2*, and *MTAP*; Figure 5) being connected to driver genes of lung adenocarcinoma (LUAD). These cases suggest that including biomarker information could enhance the effectiveness of network medicine

We found that the quality of the co-essentiality network depended on the number of cell lines whose essentiality profile was utilized to build co-essentiality links. As more cancer cell lines were compiled, co-essentiality networks were improved and identified more disease modules which can be used as potential therapeutic targets (Figure S5). Although gene essentiality data of cancer cells have been continuously accumulated in 769 cancer cell lines through various studies [65,67,78–80], the number of currently available cancer cell lines where gene essentiality profiles have been measured does not appear to be sufficiently saturated to establish complete co-essentiality links for precision oncology. Thus, the co-essentiality network has room for improvement with an increase in the number of cell lines in which essentiality profiles are currently being measured. Therefore, gene essentiality profiling of more diverse cancer cells is essential to improve the capacity of resources to define a more suitable network for precision oncology.

Like any other network-based in-silico drug repurposing method, our co-essentiality network-based method has several limitations that should be acknowledged: (1) first, while our co-essentiality network effectively prioritized anticancer targets and drug repurposing candidates, the network is based on data from cancer cell lines, which may not fully recapitulate the complexity of tumor biology in vivo. Future work could incorporate additional data from patient-derived models or in vivo studies to refine the network. (2) Second, our drug repurposing predictions were validated using clinical trial data, but further experimental validation in preclinical models would strengthen the evidence for these candidates. (3) Third, our network focuses on protein-coding genes, but non-coding RNAs and epigenetic factors also play important roles in cancer biology. Integrating these additional layers of biological complexity could further enhance the predictive power of the network. Despite these limitations, our study provides a valuable resource and framework for leveraging co-essentiality networks in precision oncology, and we anticipate that future work will build upon and extend our findings.

## Materials and methods

### Resources and network information

For all 13 networks used in this study, nodes in the networks were converted to HUGO symbols, while only the nodes and links included in the largest connected component were selected. The numbers of nodes and links of the final networks used in this study are reported in Table S2.

### Genome-wide data for network construction

#### Gene essentiality data

To build a co-essentiality network, we used a dataset consisting of the genome-wide CRISPR screenings from the Achilles project 20q2 in the dependency map (DepMap) project [19]. Specifically, the dataset was an 18,119 × 769 matrix of gene essentiality including 18,119 genes, which were screened in 769 cell lines from 26 distinct lineages using the Avana CRISPR library [81]. The dataset is publicly available at https://depmap.org/portal/download/all/ under release ‘DepMap Public 20q2’ and file ‘Achilles_gene_effect.csv’.

#### Gene expression data

To construct a co-expression network, we used CCLE expression data quantified from RNA-seq files using GTEx pipelines [82]. The dataset contains the gene expression data of 19,144 genes in 1,305 cell lines from 34 distinct lineages. Among the 19,144 genes, 23 with zero expression values across the cell lines were removed. The dataset is publicly available at https://depmap.org/portal/download/all/ under release ‘DepMap Public 20q2’ and file ‘CCLE_expression_v2.csv’.

#### Gene methylation data

To construct a co-methylation network, we used CCLE DNA methylation reduced representation bisulfite sequencing data (promoter 1 kb upstream of TSS) [82]. The dataset consists of the methylation data of 21,337 loci covering 17,182 gene promoter regions in 843 cell lines. Due to the many missing values in the data matrix, we only held cell lines with methylation data of at least 17,000 loci and loci with methylation data in at least 644 cell lines. Finally, we incorporated 20,198 methylation loci from 805 cell lines into the network construction steps. The dataset is publicly available at https://depmap.org/portal/download/all/ under release ‘CCLE 2019’ and file ‘CCLE_RRBS_TSS1kb_20181022. txt’.

### Constructing the gene correlation-based network

To construct three networks (a co-essentiality network, co-expression network, and co-methylation network) from the corresponding dataset, we measured the similarity in essentiality, expression, and methylation between two genes using corrected correlation and used this value as the link weight in the network. After data correction, we employed link filtering steps using Pearson correlation coefficient (PCC) and context likelihood relatedness (CLR) algorithm[20], as utilized in do Valle et al. [83], and filter out links that exhibited a link weight of 0. This procedure consists of three steps: (i) For all missing values in the datasets, we conducted a k-nearest neighbor (KNN) imputation with k = 10 using the impyute Python module. (ii) To measure the correlation between genes, we calculated the Pearson correlation coefficient (PCC) for all pairs of genes in the datasets and used the absolute value of PCC to capture both directions of correlation between genes. (iii) For the absolute value of PCC, we applied the context likelihood relatedness (CLR) algorithm, which conducts adaptive background correction to eliminate false correlations and indirect influences. In particular, the PCC value between gene *i* and *j*, *r_if_* was adjusted with the score 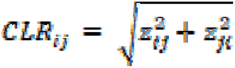 where 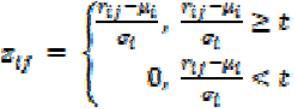, *t* is the threshold value for the *z_ij_* and *z_ij_*. The *μ_i_* and *σ_i_* are, respectively the sample mean and standard deviation of the empirical distribution of *r_ik_*, k = 1,…,n (n is the number of genes). To determine the optimal threshold t for filtering links in the three correlation-based networks (co-essentiality, co-expression, and co-methylation), we evaluated driver gene identification performance across networks constructed using six threshold values ranging from 0.0 to 5.0 (Figure S26, Table S7). We selected t = 2.0 as this threshold yielded optimal performance for co-expression and co-methylation networks in identifying cancer driver genes, while maintaining robust performance for the co-essentiality network.

One could argue that for the co-expression network, employing a non-parametric method like the Maximal Information Coefficient (MIC) might be more suitable than a parametric method such as the Pearson Correlation Coefficient (PCC), given that gene expression often deviates from normal distribution. To address this, we compared the efficacy of PCC-based and MIC-based co-expression networks in identifying cancer driver genes. At a threshold of t = 2.0, where both networks demonstrated their peak performance, the PCC-based network surpassed the MIC-based network in effectiveness (Table S8). Consequently, we opted for the PCC-based co-expression network.

### Preparation of PPI network

Eight human protein-protein interaction (PPI) networks were used: BioGRID [25], BioPlex [31], GPSnet [7], HURI [32], Inbiomap [33], iRefIndex [34], Pathway Commons [35], and STRING [36]. For all PPI networks, the values of link weights were set to 1.0.

#### BioGRID

We downloaded the BioGRID interactome from https://thebiogrid.org/ under BIOGRID-4.1.190. We used the interactions between both proteins from Homo sapiens.

#### BioPlex

We downloaded the following BioPlex interactomes from https://bioplex.hms.harvard.edu/interactions.php: BioPlex 3.0 Interactions (293T Cells) and BioPlex HCT116 v. 1.0 (HCT116 cells).We constructed a single network from the union of both interactomes.

#### GPSnet

We used the GPSnet interactome previously constructed by Cheng et al. [7], which assembled 15 commonly used databases with multiple experimental sources of evidence and an in-house systematic human protein–protein interactome. The interactome is publicly available at https://github.com/ChengF-Lab/GPSnet/tree/master/Data_mat and file ‘Net_PPI.mat’. The GPSnet was originally an *in silico* framework for drug repurposing, and in this study, we called the PPI network used in this framework GPSnet.

#### HURI

We downloaded the HURI interactome from http://www.interactome-atlas.org/ and the file ‘HuRI.tsv’.

#### InBioMap

We downloaded the InBioMap interactome from https://zs-revelen.com/download and the file ‘InBio_Map_core_2016_09_12’.

#### iRefIndex

We downloaded the iRefIndex interactome from the web interface to the Interaction Reference Index repository (iRefWeb, https://wodaklab.org/iRefWeb/) under the release iRefIndex version 13.0. We used four searching options: ‘single organism interaction‘, ‘Homo Sapiens’, ‘experimental’, and ‘physical’.

#### PathwayCommons

We downloaded the PathwayCommons interactome from http://www.pathwaycommons.org/archives/PC2/v12/ and the file ‘PathwayCommons12.All.hgnc.txt’.

#### STRING

We downloaded the STRING interactome from https://string-db.org/ under the release v11.0. To avoid co-citation information in STRING, we removed the text-mining scores for all links and recalculated the confidence score of STRING. Next, links with confidence scores > 700 were considered to leverage the high-confidence PPIs.

### Benchmark co-essentiality networks

Four co-essentiality networks from three published studies were used as benchmarks for our approaches. We reconstructed each network using the same gene essentiality dataset employed in our co-essentiality network construction (18,119 genes × 769 cell lines) to eliminate performance bias from different cell line compositions. The first benchmark network, Wainberg_etal [11], was built using generalized least squares correlation. The second network, Amici_etal [12], was constructed using a rank-based bottom-up approach with a rank threshold of 30, selected in the original paper. The remaining two networks from Gheorghe et al. [37] used the PCA whitening method in that study but differed in their essentiality measurements: Gheorghe_etal_Ceres utilized pre-calculated Ceres scores from DepMap, while Gheorghe_etal_BF employed Bayes Factors calculated using BAGEL2. The networks were constructed following each study’s methods and source code.

### Genetic interaction network

To compare the co-essentiality network with the genetic interaction network based on a synthetic-lethal relationship, we used a clinically relevant synthetic lethality network built by the ‘identification of clinically relevant synthetic lethality (ISLE)’ approach [84]. We downloaded ‘clinically relevant synthetic lethality network (cSLnet)’ interactome from https://github.com/jooslee/ISLE/tree/main/networks and the file ‘ISLE_clinical_SL_network_FDR_0.2.cys’.

### Essential gene fraction in the networks

To measure fraction of essential genes in the network, we used the 2,123 common essential genes in the DepMap. The dataset is publicly available at https://depmap.org/portal/download/all/ under release ‘DepMap Public 20q2’ and file ‘Achilles_common_essentials.csv.

### Curated gene set enrichment analysis (GSEA) of co-essentiality network

We calculated the enrichment of co-essentiality links in six curated gene sets. We downloaded six curated gene sets: molecular pathways (KEGG [21], REACTOME [23]) and Gene Ontology annotations [24] [GO: BP (Biological Process), MF (Molecular Function), and CC (Cellular Components)] from Molecular signatures database(MSigDB) [85], and human core protein complex from CORUM [22].

In the co-essentiality network, gene pairs were ranked by the link weights and grouped into cumulative bins of 10,000 pairs, and the enrichment was calculated using the ratio of pairs annotated with the same biological modules. We also measured enrichment expected by chance as the probability of finding the gene pairs within the same biological modules without being informed by co-essentiality links. Similar to the approach of Lee et al. [86], for the expected ratio, we changed the denominator from the number of co-essentiality links to all possible pairs between the genes given a bin.

### Modularity calculation

We used two types of modularity measures as shown in the following function: 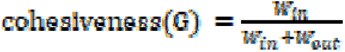, where *W_in_* is the total weight of links exclusively present within group of genes G, *W_out_* is the total weight of links that connect the gene group with the rest of the network [87] and 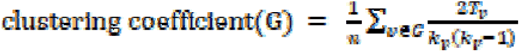, where *n* is the number of the gene group G, *T_v_* is the number of triangles through node, and *k_v_* is the degree of node v. Since both modularity measures can be affected by the degree centrality of genes in a network, we applied normalization to the modularity measures, which removes the degree bias. Similar to the approach of Guney et al. [54], we created a reference modularity distribution that corresponds to the expected modularity of 100 randomly selected groups of genes matching the size and degrees of query genes in the network. Next, modularity was normalized as the z-score of the observed modularity of genes computed from the reference distribution of modularity from random groups.

### Network enrichment to cancer related pathways (CRPs)

Among the 186 KEGG pathways from MSigDB, we used 31 as CRPs, which included the related pathways of ‘Pathways in cancer’(Pathway ID: hsa05200) and the pathways in ‘Cancer: specific types’. A list of the 31 KEGG pathways is shown in Table S1.

For 186 KEGG pathways, we calculated relative modularity by converting scaled modularity calculated using cohesiveness to rank percentile scores in four networks: the co-essentiality network, PPI network (BioGRID), co-expression network, and co-methylation network. Next, we defined network-specific pathways by assigning the type of network to each of the 186 KEGG pathways, according to which network showed the highest relative modularity. The relative modularity values are presented in Table S1.

Finally, for each network, we constructed a 2×2 contingency table with four types of pathways namely network-specific CRPs, non-network-specific CRPs, network-specific non-CRPs, and non-network-specific non-CRPs. From the contingency table, we calculated the odds ratio of network enrichment to CRPs and determined the statistical significance of enrichment by calculating the p-value from Fisher’s exact test. The contingency table of the co-essentiality network is shown in the Fig 1C.

### Driver genes of each cancer type

#### Bailey et al

We retrieved driver genes of each cancer type from Bailey et al. [88], who reported 299 driver genes across 33 cancer types. For modularity analysis, we selected 19 cancer types with more than 10 reported driver genes included in the co-essentiality network: bladder urothelial carcinoma (BLCA), breast invasive carcinoma (BRCA), cervical squamous cell carcinoma and endocervical adenocarcinoma (CESC), Colorectal adenocarcinoma (COADREAD), lymphoid neoplasm diffuse large B-cell lymphoma (DLBC), esophageal carcinoma (ESCA), glioblastoma multiforme (GBM), head and neck squamous cell carcinoma (HNSC), kidney renal clear cell carcinoma (KIRC), acute myeloid leukemia (LAML), brain lower grade glioma (LGG), liver hepatocellular carcinoma(LIHC), lung adenocarcinoma (LUAD), lung squamous cell carcinoma (LUSC), prostate adenocarcinoma (PRAD), skin cutaneous melanoma (SKCM), stomach adenocarcinoma (STAD), Uterine Corpus Endometrial carcinoma (UCEC), and uterine Carcinosarcoma (UCS).

#### Cancer Gene Census (CGC)

For further validation of the high modularity of driver genes in the co-essentiality network, we used the experimentally validated cancer driver gene set from Cancer Gene Census (CGC) database [30]. For cancer type specific analysis, we manually mapped Tier 1 driver genes in CGC to TCGA cancer types according to their tumor type information. A total of 470 driver genes were mapped to 22 TCGA cancer types. Similarly, for modularity analysis, we selected 19 cancer types with more than 10 reported driver genes included in the co-essentiality network: bladder urothelial carcinoma (BLCA), breast invasive carcinoma (BRCA), Colon adenocarcinoma (COAD), lymphoid neoplasm diffuse large B-cell lymphoma (DLBC), head and neck squamous cell carcinoma (HNSC), Kidney Chromophobe (KICH), kidney renal clear cell carcinoma (KIRC), acute myeloid leukemia (LAML), brain lower grade glioma (LGG), liver hepatocellular carcinoma (LIHC), lung adenocarcinoma (LUAD), lung squamous cell carcinoma (LUSC), Ovarian serous cystadenocarcinoma (OV), Pancreatic adenocarcinoma (PAAD), prostate adenocarcinoma (PRAD), skin cutaneous melanoma (SKCM), stomach adenocarcinoma (STAD), Thyroid carcinoma (THCA), and Uterine Corpus Endometrial carcinoma (UCEC). The driver genes mapped to cancer types are listed in Table S9.

### Driver gene prediction with guilt-by-association

To evaluate the performance of each network for driver gene identification, we conducted leave-one-out cross validation with driver genes from Bailey et al. [88]. For each network, we measured the prediction score of each gene via the ratio of the number of driver genes in the first neighbors of the gene as follows: 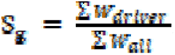, where S_g_ is the prediction score of gene g, *W_driver_* is the weight of links from gene g connected to driver genes, and *W_all_* is the weight of all links from gene g. The performance of the network was measured using the area under the curve of the receiver operating characteristic curve (AUROC) on the prediction score, which was designed to obtain a high value when driver genes were assigned a high prediction score.

For the ‘without-network’ control method, we calculated the prediction score using the average gene essentiality value of cell lines within the same cancer type. Using this prediction score, we measured the performance of driver gene identification using the same AUROC metric as applied in network-based approaches.

### Driver gene prediction with Hotnet2

To evaluate the performance of driver gene identification for each network, we also used a network propagation method, Hotnet2 [38], which is an algorithm to identify significantly mutated subnetworks [38]. As inputs of HotNet2, we employed (i) the mutation data of 11 cancer types used in the original paper of HotNet2 and (ii) the networks used in this study, including the co-essentiality network. The mutation data were preprocessed using the same process described in the original paper. The parameters for the permutation numbers were set to 100 for the networks and 1,000 for the heat. The insulating parameter β was set to 0.04. The default settings for other parameters were used. The source code of Hotnet2 algorithm of the original authors[38] at a GitHub repository (https://github.com/raphael-group/hotnet2) was used.

The performance of Hotnet2 was measured with the number of driver genes included in the modules resulting from Hotnet2 using the F1 score. For each cancer type, the driver genes from Bailey et al.[88] were used as a positive set of predictions. The F1 score was calculated as follows:
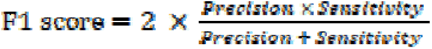

### Driver gene prediction with uKIN

To test the performance of driver gene identification for each network, we used a guided network propagation method, uKIN [39], which uses prior knowledge of cancer genes to guide, within networks, the network propagation that starts from newly identified candidate genes. We built an in-house script for the uKIN algorithm, which is available at https://github.com/SBIlab/CESnet-repurposing.

Adapting the guided propagation concepts from uKIN, we conducted two stages of network propagation using the page-rank algorithm from the NetworkX Python module [89]. In the first stage, we used one for genes from prior knowledge in the network and zero for all other genes in the network as an input for the personalization parameter in the page-rank algorithm. From the results of the network propagation, we redevised the network as a directed graph and re-assigned link weights as follows:

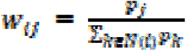

where *w_ij_* is the weight of the link from node i to node j, *p_k_* is the propagation score of node k, and *N*(*t*) are the neighbors of node i. In the second stage, we conducted network propagation with a redevised network, using new information as seeds. All other page-rank algorithm parameters were adjusted to their default values (damping factor = 0.85).

The performance of the guided network propagation was evaluated using the same method used in the uKIN study. We used 723 cancer gene census (CGC) [30] genes as the prior and positive sets, respectively. First, we separated the CGC genes that occur in our network into two categories, namely prior knowledge set and hidden set, which indicate seed genes for the first propagation and test set for the ukin outcome, respectively. To create the group of positives that we hoped to find, we randomly selected 400 genes from the CGC as a hidden set for evaluation. From the remaining 323 CGCs, we randomly selected 20 genes from the network as prior knowledge. The other CGC genes that were not included in the prior knowledge set or hidden set were discarded. Different hidden sets and prior knowledge sets were used for 100 iterations of the uKIN algorithm. After the first propagation, the mutational frequency of each gene was used as the input for the second propagation. We utilized somatic mutation data of 24 cancer types used in the uKIN study as new information inputs of uKIN obtained from Tokheim et al. [90] who collected 1,225,917 somatic mutations in 8,657 samples of TCGA somatic mutation calls (v0.2.8, https://synapse.org/MC3) [91]. We calculated the mutational frequency of a gene by dividing the frequency of somatic missense and nonsense mutations found in tumor samples by the length of the protein. To evaluate the performance of driver gene prediction, we calculated the AUROC of the top 100 scored genes resulting from the guided propagation. To compute the AUROC, only the CGC genes included in the hidden set were regarded as positive, and the other genes in the network were considered negative. Finally, we calculated the mean value from the AUROC from the 100 iterations.

### Network propagation with driver genes of each cancer type

To prioritize genes in the network according to their distance from the driver gene, we conducted network propagation using the page-rank algorithm from the NetworkX Python module [89]. Among the 19 cancer types, we conducted network propagation for 17 cancer types, where the co-essentiality network had the highest modularity. For each cancer type, we assigned one for driver genes and zero for other genes in the network as input for the personalization parameter in the page-rank algorithm. All other page-rank algorithm parameters were adjusted to their default values (damping factor = 0.85).

### Patient stratification and survival analysis

To identify driver modules capable of differentiating patient survival, we identified biological pathways located proximal to driver genes using network propagation scores of the genes in a network. To calculate the propagation score of the genes included in each pathway, we conducted a GSEA using the GSEApy Python module [92]. Through GSEA on network propagation score, we selected pathways significantly enriched in genes with high network propagation scores using an FDR of < 0.001 and NES of > 0 as driver modules. On the other hand, the ‘without-network’ control method used only the driver genes of each cancer type as a driver module.

The transcriptome and clinical tables of patients used in this study were downloaded using the TCGAbiolinks R package [93]. For the pre-processing of gene expression data of TCGA patients, we computed the gene expression levels using read counts, which were normalized by gene length corrected trimmed mean of M-values [94] calculated using the edgeR [95] R package. For statistical significance, we selected cancer types that contained at least 100 samples with survival data among 17 cancer types, where the co-essentiality network had the highest modularity of the driver genes. This resulted in 16 cancer types, including 7,259 tumor samples.

We stratified the patients from TCGA into two groups according to their gene expression levels in the driver modules. We conducted single-sample gene set enrichment analysis (ssGSEA) on the gene expression of the selected driver modules for each patient. According to NES of each patient, we defined the top 50% of patients as the upregulated group and the bottom 50% of patients as the downregulated group. To determine whether a survival difference existed between the upregulated and downregulated groups, we used the log-rank test and obtained statistical significance. To compare the maximum capability of patient subtyping across the networks, the driver module from each network whose patient grouping had the lowest p-value from the log-rank test was selected, and its p-value is shown in Figure 2I.

### Drug-gene association data

We obtained drug-gene association data from PanDrugs [53] which assembled 18 resources with data curated by experts and drug-gene associations collected from experimental drug screenings. The source data of PanDrugs was provided by the original authors. We used two drug-gene association types, “direct target” and “biomarker”, as drug targets. From the 9,090 drugs listed in PanDrugs, only those with a PubChem ID were selected to prevent duplicates caused by synonym usage. Finally, we considered 35,486 drug-gene associations across 7,015 drugs (Table S10).

### Performance of prioritizing approved anticancer DAGs

To determine whether network propagation results can prioritize the DAGs of FDA-approved drugs in each cancer type, we performed GSEA on the propagation results of driver genes from the query cancer type using a DAG list of approved drugs as a gene set. We conducted a GSEA using the GSEApy Python module [92]. We collected FDA-approved drugs across 16 cancer types from the TCGA(https://www.cancer.gov/about-cancer/treatment/drugs) and PanDrugs database. The synonyms of drugs between TCGA and drug-gene association data were mapped with PubChem ID using the Pubchempy Python module [96]. There is no FDA approved drugs of ESCA mapped to drug-gene association data, we excluded ESCA for this analysis. Finally, 176 FDA approved drugs across the 16 cancer types were selected for the analysis. For each cancer type, a list of genes associated with approved drugs was used for GSEA. The drugs’ annotation between TCGA and Pandrugs were mapped by PubChem ID. The performance of the network in prioritizing approved anticancer DAGs was measured using the NES value of the GSEA results. The FDA-approved anticancer drugs and their DAGs used in this analysis are listed in Table S3.

For the ‘without-network’ control method, we conducted GSEA using the average gene essentiality value of cell lines within the same cancer type.

### Calculation of TC score

For 17 cancer types, we assigned a therapeutic candidate (TC) for each drug. The TC score is aggregated from the network propagation value of the DAGs by taking their root-mean-square (RMS) as follows:

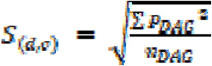

where *S*(*_d,v_*) is the TC score of the drug d in cancer type c, *P_DAG_* is the propagation score of the DAGs in cancer type c and *n_DAG_* is the number of DAGs.

### Performance of predicting the cytotoxic effect of drugs

To obtain drug response data of human cancer cell-line, we downloaded ‘secondary-screen-dose-response-curve-parameters’ from PRISM database [97], which included 1,448 compounds screened against 499 cell lines. Among the 17 cancer types, we used 15 cancer types, including cancer cell lines, which have secondary screen data in PRISM. The cytotoxic effect of the drug on each cancer type was calculated by taking the median IC_50_ values of the drug in cancer cell lines belonging to the corresponding cancer.

The network’s performance in predicting the cytotoxic effect of drugs on cancer was measured using the Spearman correlation coefficient between TC score and negative log base ten of the drug’s median IC_50_ values for the cancer. For the ‘without-network’ control method, instead of the TC score, we used the number of driver genes among DAGs of each drug as the prediction score. The synonyms of drugs between PRSIM and PanDrugs were mapped with PubChem ID using the Pubchempy Python module [96].

### Performance of predicting reversal gene expression(RGE) effect of drugs in COADREAD

To estimate the efficacy of the drug by altering the gene expression to be a reversal of that in cancer conditions, we used the L1000 dataset from CMap[52]. From the L1000 dataset, we obtained 6,056 compounds with PubChem IDs and expression profiles for 978 genes directly measured in 15 cancer cell lines whose primary site was the large intestine: CL34, HCT116, HELA, HT115, HT29, LOVO, MDST8, NCIH508, NCIH716, RKO, SNU1040, SNUC5, SW480, SW620, and SW948. We initially computed the RGE effect between the differential gene expression of COADREAD (obtained via CREEDS [51], signature ID: dz552) and the compound’s transcriptional profile using Zhang’s connection score [50], that is, the absolute value of the anti-correlated, standardized ranked connection score. Since drugs have multiple transcriptional profiles for each cell line depending on the treatment dose and time, we calculated the max RGE effect for each cell line and used the median RGE effect across cell lines as a representative value. Finally, the performance of the network in predicting the drug’s RGE effect was measured using the Spearman correlation coefficient between the TC score and the RGE effect of drugs that exist in both PanDrugs and CMap.

### Selection of repurposing candidates

Among the 1,213 drugs approved in non-cancer disease with drug target information in PanDrugs [53], we measured the significance of the TC score by empirical p-value calculated from permutation test of drug-associated genes(DAGs). Similar to the approach used for normalized modularity calculation, we constructed a reference TC score distribution for each drug. This distribution corresponded to the expected TC score of 100 randomly selected groups of genes that matched both the size and degrees of the DAGs within the network (see Figure S21). To assess the significance of the TC score, we calculated its p-value based on the reference distribution, which was derived from random DAGs. Subsequently, we applied the Benjamini-Hochberg procedure for multiple test correction to the p-values, and the resulting adjusted P-values were utilized to determine the significance of the TC score.

We selected drugs with significant TC score (adjusted P <= 0.05) as repurposing candidates. This resulted in 145 approved drugs with new therapeutic indications across the 17 cancer types (Table S5).

### Performance of prioritizing drug repurposing candidates

To validate the repurposing candidates identified through the co-essentiality network, we leveraged clinical trial information. We considered 169 drugs labeled as “CLINICAL_CANCER,” indicating their presence in clinical trial records related to cancer, as a positive set for drug repurposing (Table S10). We assessed the performance of the drug repurposing by measuring the number of repurposing candidates (with adjusted P-values ≤ 0.05) that were included in the positive set, using the F1 score as our evaluation metric.

### Comparative analysis with an existing network-based drug repurposing method

To validate the performance of drug repurposing, we implemented the network proximity-based approach presented in Cheng et al.[16]. For each cancer type, the proximity between cancer driver genes and drug-associated genes (DAGs) was calculated using the shortest path distances in the network. The proximity score d(S,T) between a set of cancer driver genes (S) and DAGs (T) was defined as:

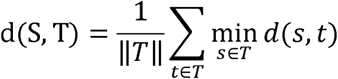

where d(s,t) is the shortest path length between nodes s and t. The proximity scores were converted to z-scores by comparing them against a reference distribution generated from 1000 randomly selected gene sets matched in size and degree to the original sets. In addition to evaluating the original z-score threshold of z < −0.15, we also assessed drug repurposing candidates using two less stringent thresholds (z < −0.1 and z < −0.05) to ensure robust comparison. The performance was evaluated using drugs with cancer-related clinical trials as the gold standard. The toolbox package for the network proximity calculation can be downloaded at github.com/emreg00/toolbox. The human interactome used to measure network proximity was downloaded from the original paper[16], Supplementary Data 1.

## Code availability

The source codes for reproduction of the results were developed in either python 2.7.13 or python 3.7.9 and are available at a GitHub repository(https://github.com/SBIlab/CESnet-repurposing).

## Data availability

All data are available in the main text, the supplementary materials or public repository. The final 16 networks used for the analysis in this study are available at the Mendeley Data repository (https://data.mendeley.com/preview/v5ydnp4p9j?a=43e34493-78cb-420b-9008-73acd8dbb9fb).

## Authors’ contributions

**KL:** Conceptualization, Data curation, Investigation, Writing - original draft, Visualization, Investigation, Validation, Methodology, Project administration. **DK:** Investigation, Writing - original draft, Investigation. **IK:** Conceptualization, Writing - original draft, Methodology. **JL:** Investigation, Methodology. **DH:** Methodology, Writing - original draft. **SL:** Data curation, Investigation. **EK:** Methodology. **SI:** Conceptualization, Supervision. **KS:** Conceptualization, Supervision. **SK:** Conceptualization, Funding acquisition, Supervision, Project administration, Writing - original draft. All authors read and approved the final manuscript.

## Acknowledgments

We thank all members of the Kim laboratory for their helpful discussions. We would also like to thank Pineiro Elena and Professor Al-Shahrour. Fatima for providing the PanDrugs dataset. This work was supported by grants from the Korean National Research Foundation (2021R1A2B5B01001903 and 2020R1A6A1A03047902) and IITP (2019-0-01906, Artificial Intelligence Graduate School Program, POSTECH).

