## Supplemental_materials for "A Co-essentiality Network of Cancer Driver Genes Better Prioritizes Anticancer Drugs"

**Supplemental material for “A co-essentiality network of driver genes better prioritizes anticancer drugs**

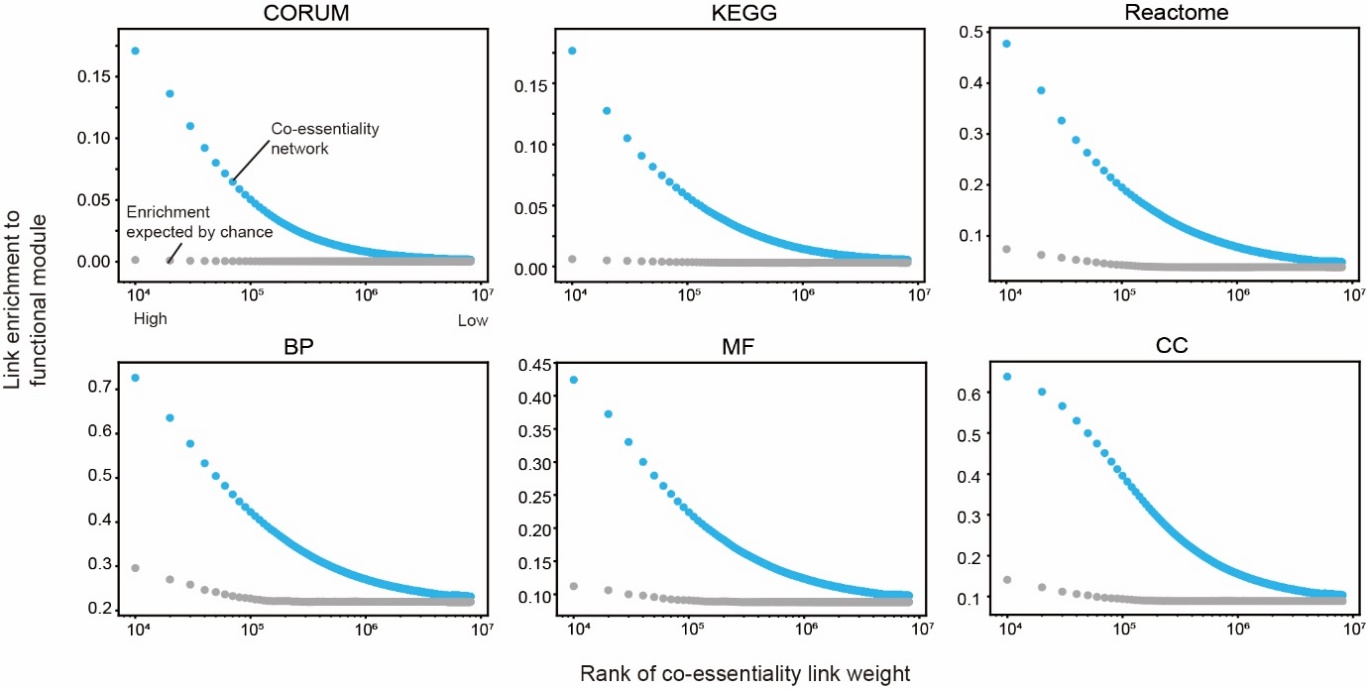

**Figure S1. Curated gene set enrichment of co-essentiality links.** Enrichment of the co-essentiality links to six curated gene sets: CORUM, KEGG, Reactome, GO:BP, GO:MF, and GO:CC. Co-essentiality links were ranked and binned (n = 10,000). The enrichment of the co-essentiality links in each bin is indicated in blue dots. The expected enrichment of co-essentiality links in each bin is indicated by gray dots.

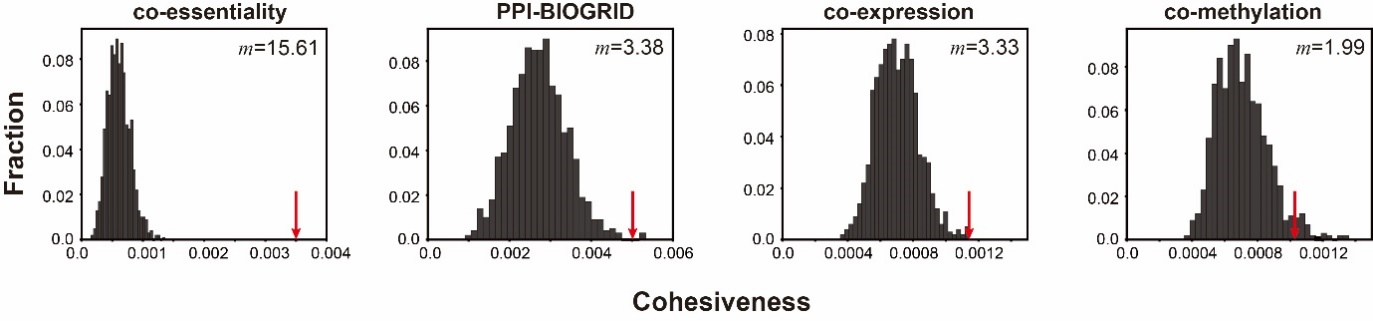

**Figure S2. Empirical distribution of modularity of the degree controlled random nodes for LUSC driver genes.** The x-axis showed the value of cohesiveness used for the modularity measure, while the y-axis denotes the fraction of the number of random modules against 100 permutations. The red pointers are observed cohesiveness value from each network. The value *m* in the upper right corner is the normalized modularity value calculated from the empirical distribution of the cohesiveness values.

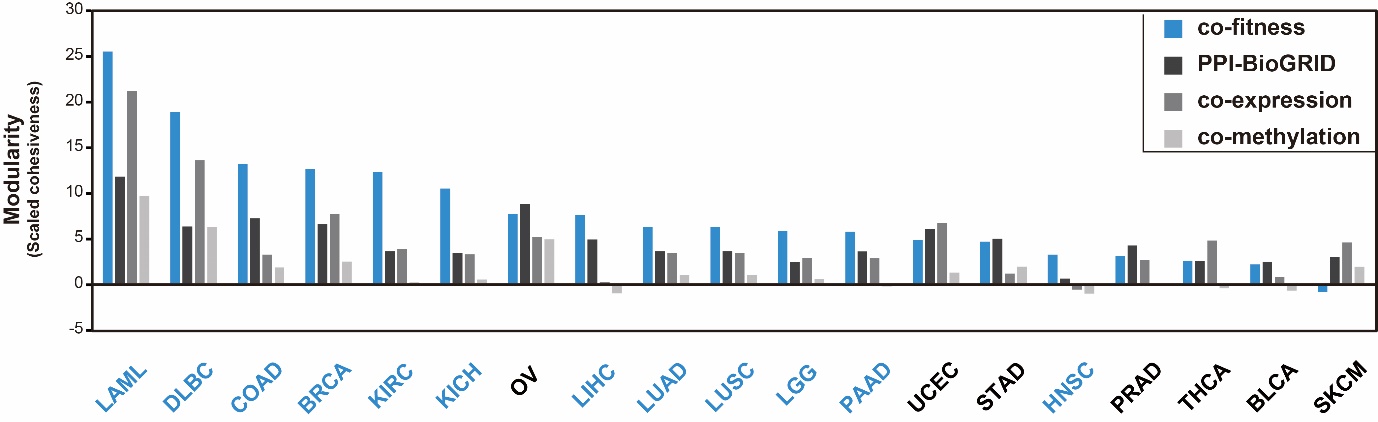

**Figure S3. Modularity of CGC cancer drivers in the four networks.** The modularity calculated from the subnetwork of driver genes from CGC for the four networks: co-essentiality, PPI-BioGRID, co-expression, and co-methylation. For blue-colored cancer types, the co-essentiality network showed the highest modularity among the four networks.

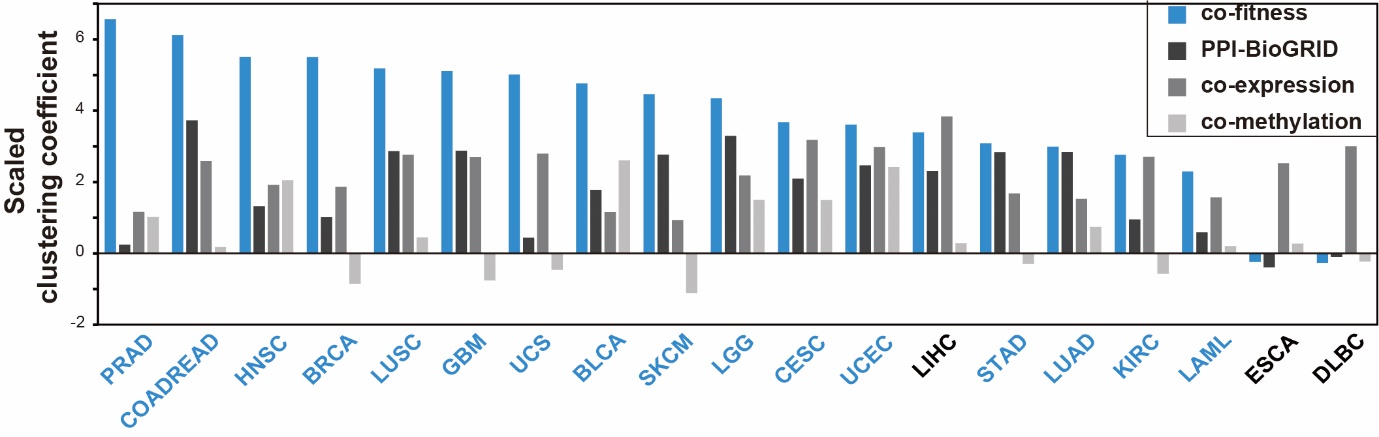

**Figure S4. Scaled clustering coefficient of cancer driver genes in the four networks.** Scaled clustering coefficient calculated from the subnetwork of driver genes across 19 TCGA cancer types for the four networks: co-essentiality, PPI-BioGRID, co-expression, and co-methylation. For blue-colored cancer types, the co-essentiality network showed the highest scaled clustering coefficient among the four networks.

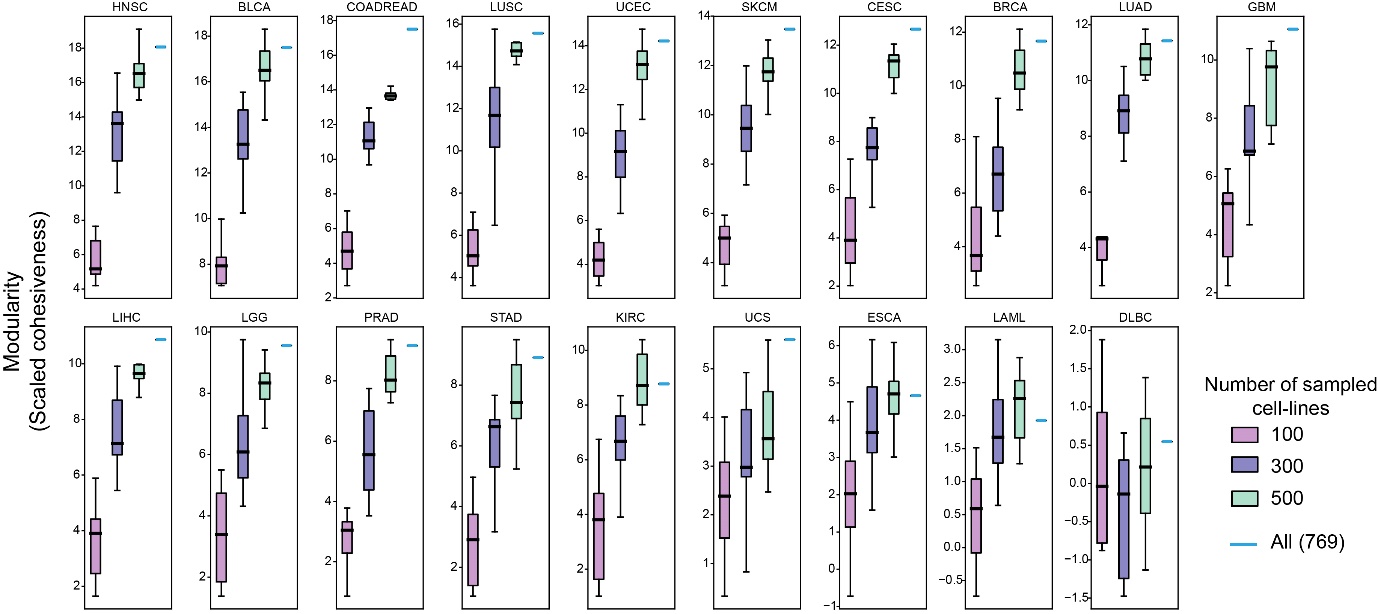

**Figure S5. Correlation between the modularity of driver genes in the co-essentiality network and the number of cell lines used to construct the co-essentiality network.** The purple, blue, and green colors represent co-essentiality networks constructed with 100, 300, and 500 randomly selected cell lines, respectively. The sky-blue lines indicate the co-essentiality network constructed using all 769 cell lines in the DepMap 20q2 dataset.

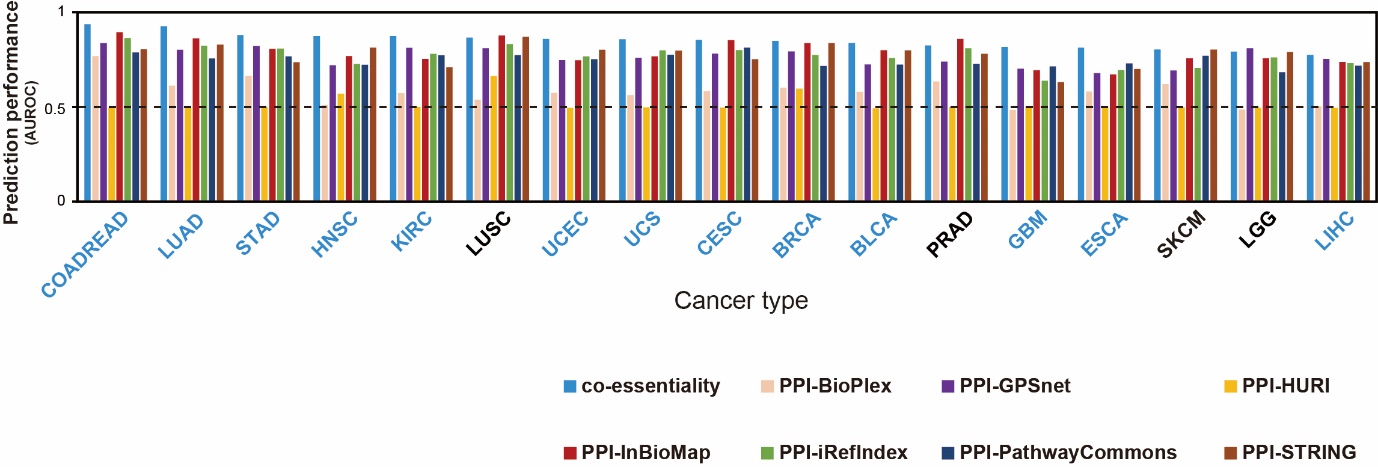

**Figure S6. Performance of the co-essentiality network for cancer driver gene identification compared with that of seven different PPI networks.** Performance of driver gene identification for eight networks: co-essentiality, BioPlex, GPSnet, HURI, Inbiomap, iRefIndex, Pathway Commons, and STRING. For blue-colored cancer types, the co-essentiality network showed the best performance among the eight networks.

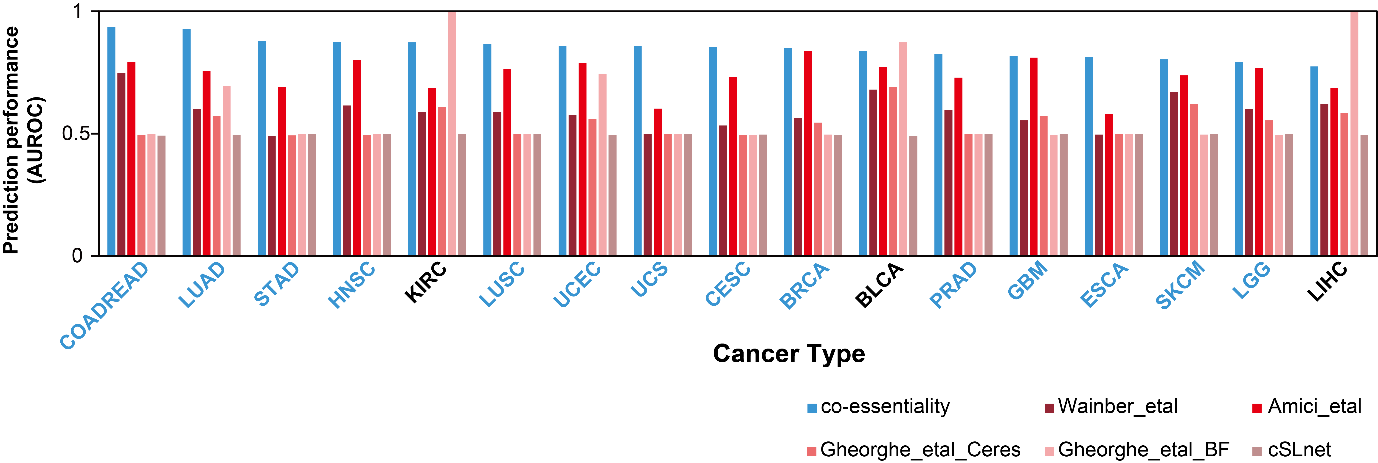

**Figure S7. Performance of the co-essentiality network for cancer driver gene identification compared with that of other co-essentiality networks and the genetic interaction network.** Performance of driver gene identification for other co-essentiality networks (Wainberg_etal, Amici_etal, Gheorghe_etal_Ceres, and Gheorghe_etal_BF), and the genetic interaction network based on synthetic-lethal relationship (cSLnet). For blue-colored cancer types, the co-essentiality network showed the best performance among the three networks.

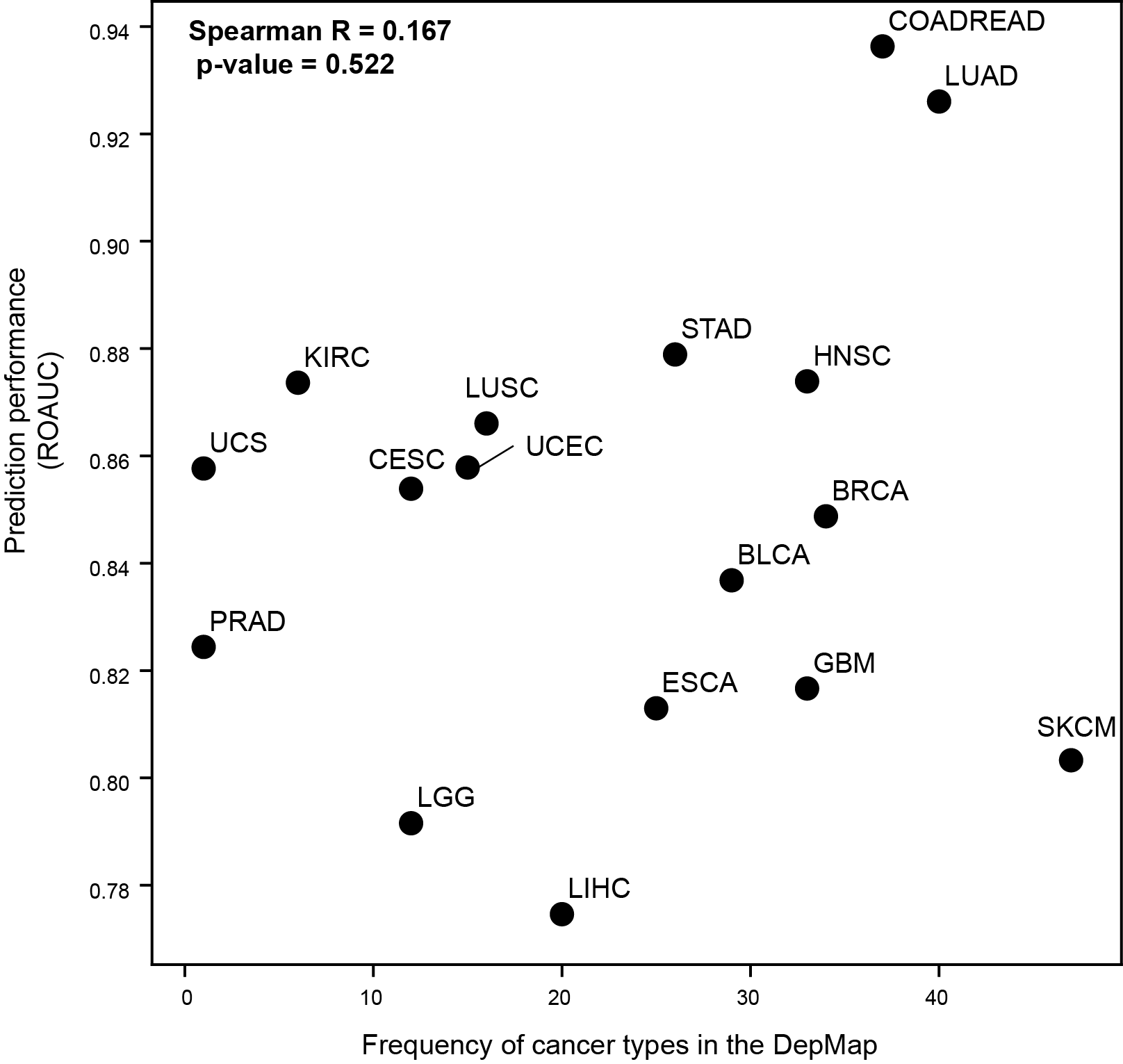

**Figure S8. Correlation between performance of the co-essentiality network for cancer driver gene identification and frequency of cancer types used for co-essentiality network construction.** The correlation was calculated by Spearman correlation coefficient R.

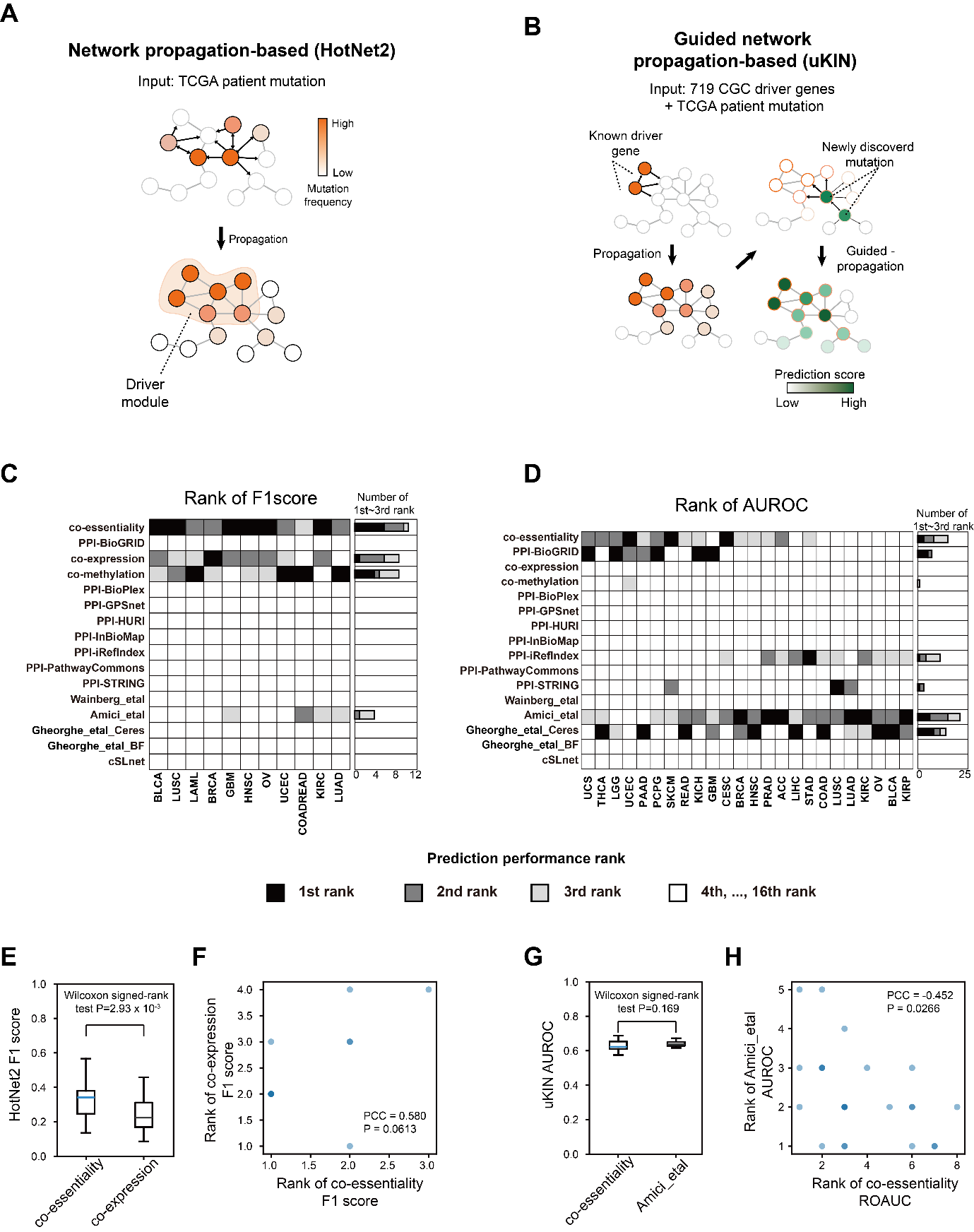

**Figure S9. Performance for Hotnet2-based and uKIN-based driver gene identification. (A, B)** Schematic overviews of the HotNet2 and uKIN algorithms. **(C, D)** Heatmaps showing performance ranks of driver gene identification across 16 networks, using HotNet2 F1 scores **(C)** and uKIN AUROC **(D)**. Black, dark gray, and light gray indicate 1st, 2nd, and 3rd place, respectively; white indicates ranks below 4th. **(E, G)** Boxplots comparing the top two performing networks for HotNet2 F1 scores (**E**: co-essentiality vs. co-expression) and uKIN AUROC (**G**: co-essentiality vs. Amici_etal). Significance was assessed by the Wilcoxon signed-rank test**. (F, H)** Scatter plots comparing performance ranks across cancer types for HotNet2 F1 scores (F, 11 cancer types; x-axis: the co-essentiality network, y-axis: the co-expression network) and uKIN AUROC (**H**, 24 cancer types; x-axis: the co-essentiality network, y-axis: Amici_etal). Pearson’s correlation coefficient was used to measure correlation between the two network’s ranks.

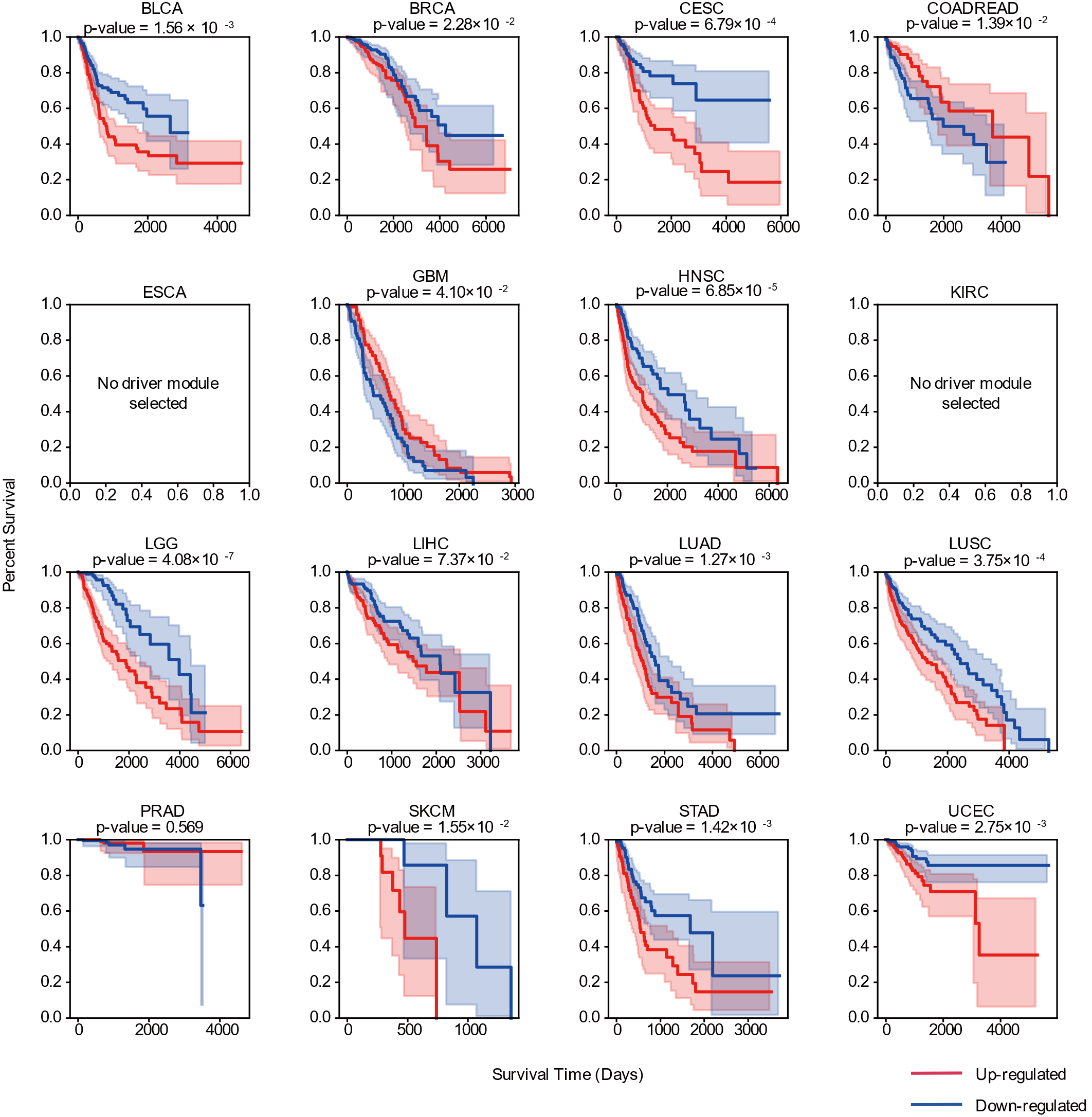

**Figure S10. Survival plot of patient subgroups in 16 cancer types stratified by the co-essentiality network.** Red lines represent patient groups with upregulated driver module expression. Blue lines represent patient groups with downregulated driver module expression. Significance of survival difference between patient groups is shown as the p-value of log-rank test. Cancer types denoted with a blank box had no significant driver modules in the network.

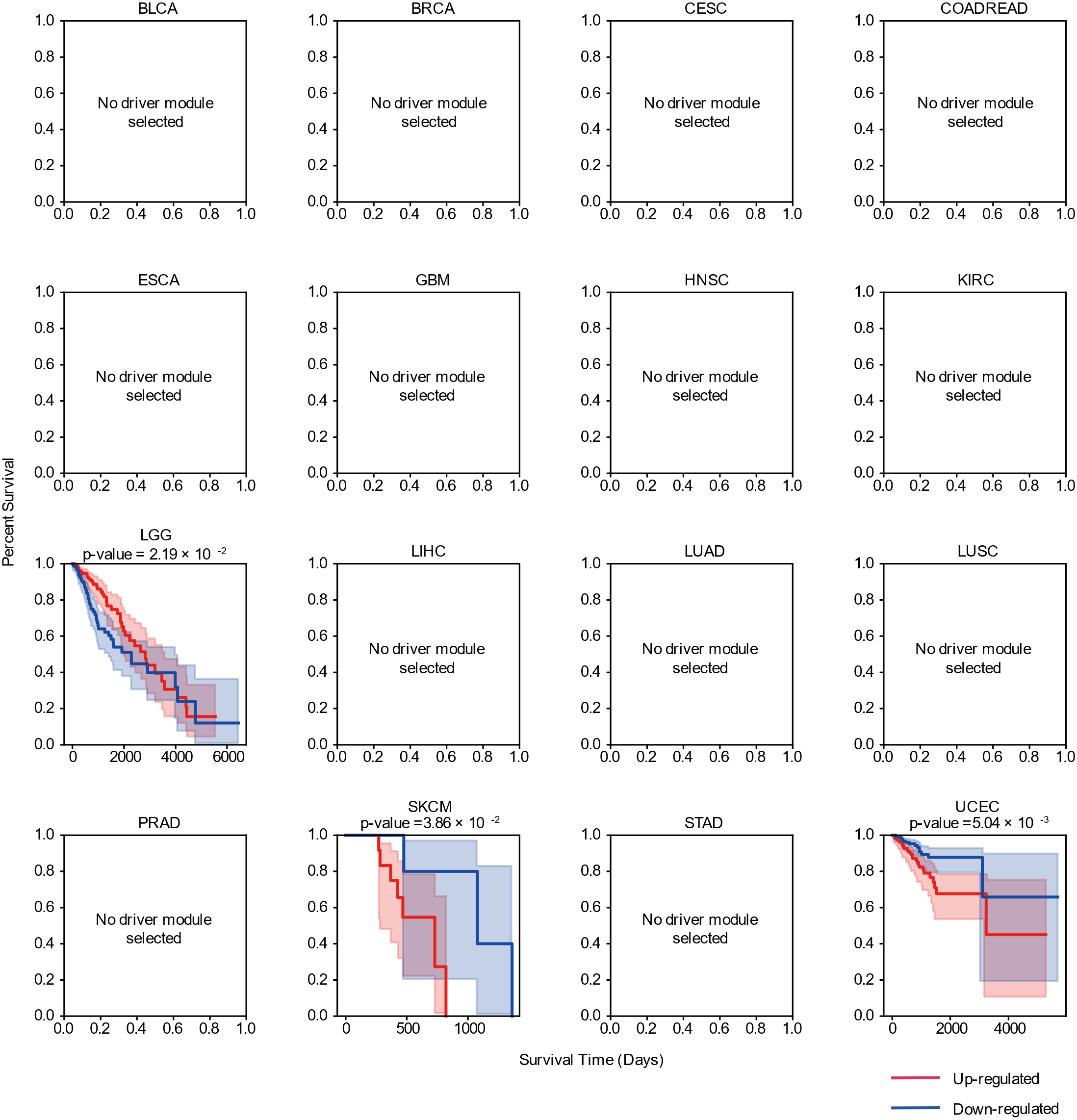

**Figure S11. Survival plot of patient subgroups in 16 cancer types stratified by the PPI-BioGRID.** Red lines represent patient groups with upregulated driver module expression. Blue lines represent patient groups with downregulated driver module expression. Significance of survival difference between patient groups is shown as the p-value of log-rank test. Cancer types denoted with a blank box had no significant driver modules in the network.

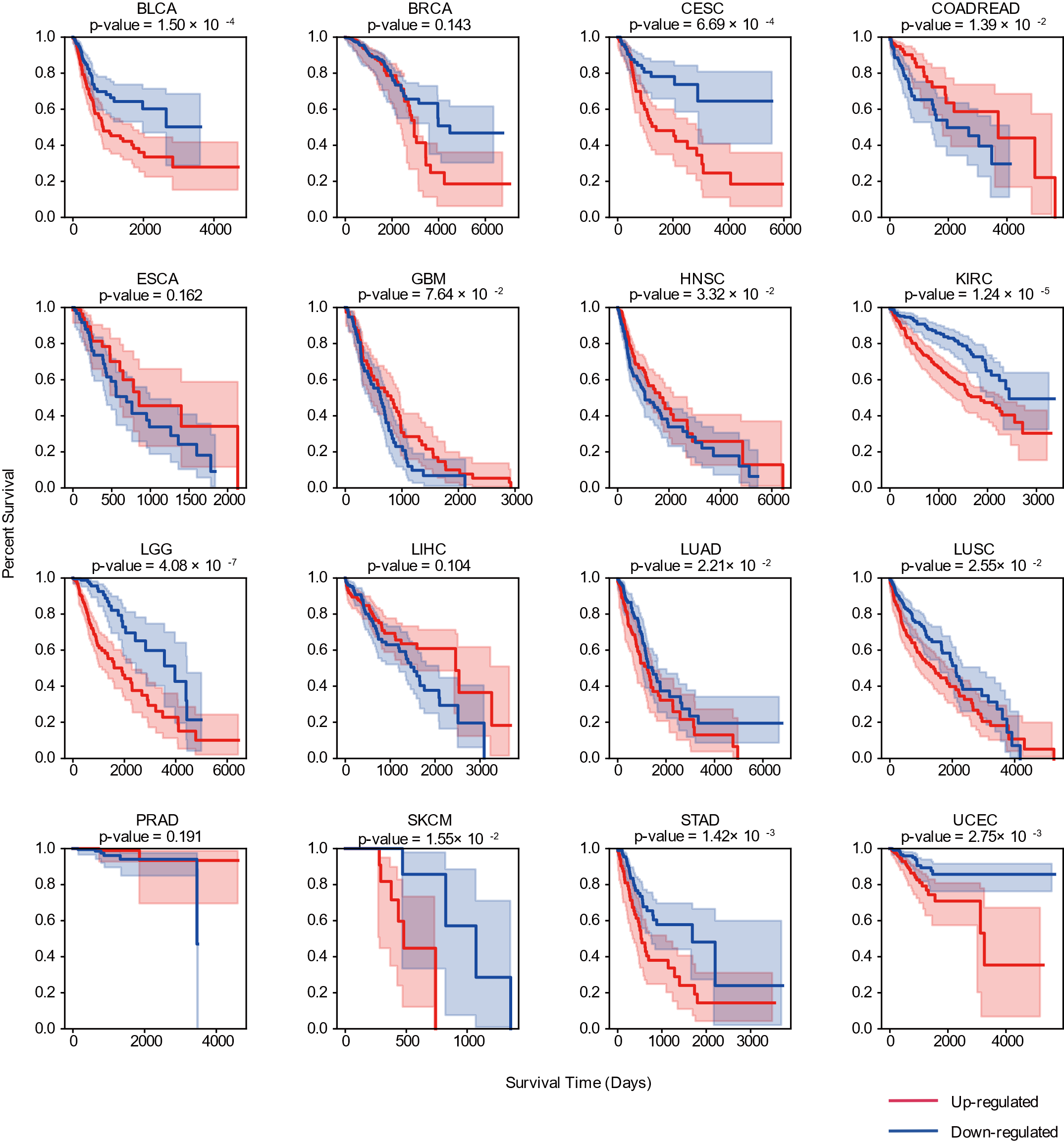

**Figure S12. Survival plot of patient subgroups in 16 cancer types stratified by the co-expression network.** Red lines represent patient groups with upregulated driver module expression. Blue lines represent patient groups with downregulated driver module expression. Significance of survival difference between patient groups is shown as the p-value of log-rank test. Cancer types denoted with a blank box had no significant driver modules in the network.

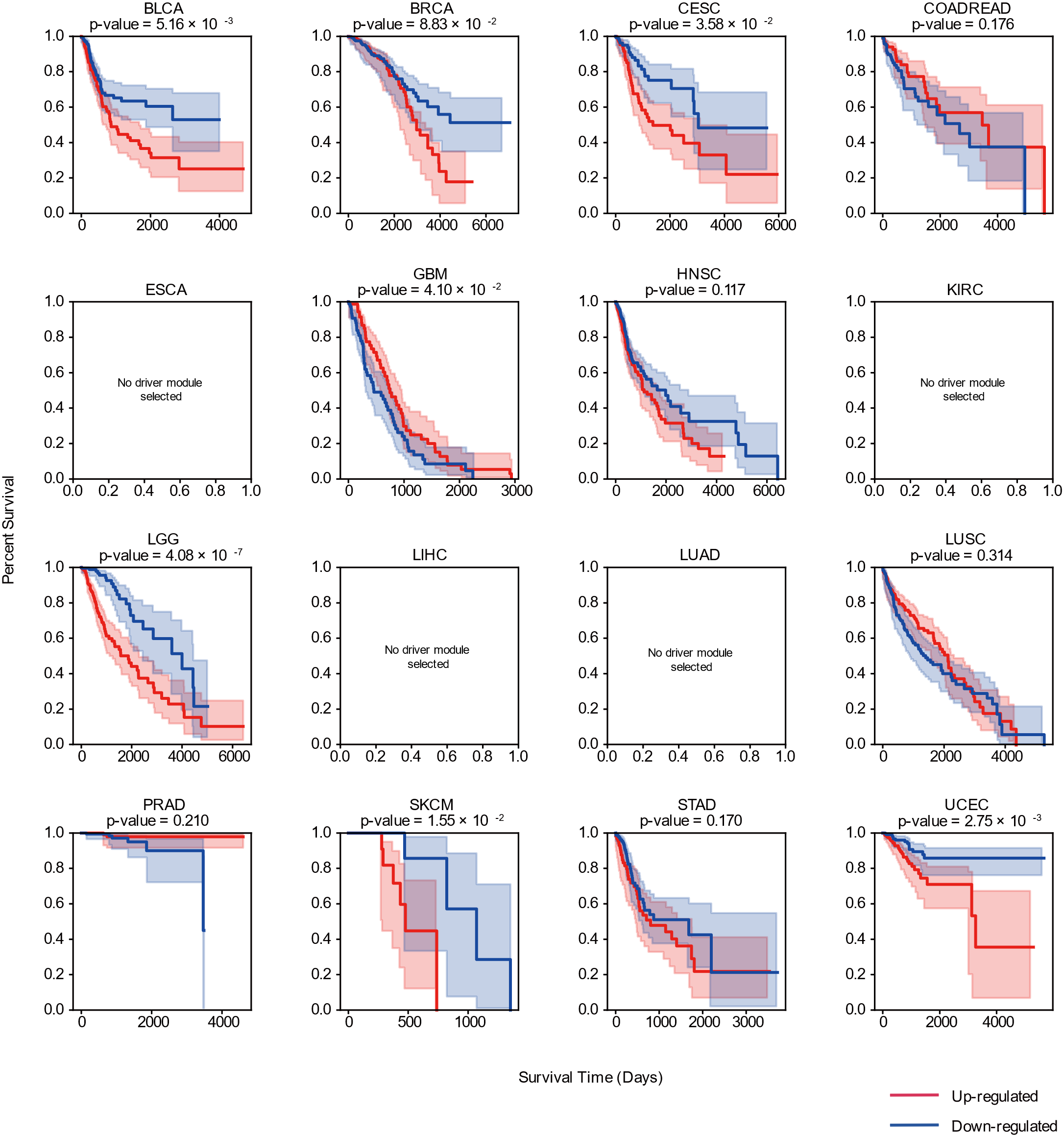

**Figure S13. Survival plot of patient subgroups in 16 cancer types stratified by the co-methylation network.** Red lines represent patient groups with upregulated driver module expression. Blue lines represent patient groups with downregulated driver module expression. Significance of survival difference between patient groups is shown as the p-value of log-rank test. Cancer types denoted with a blank box had no significant driver modules in the network.

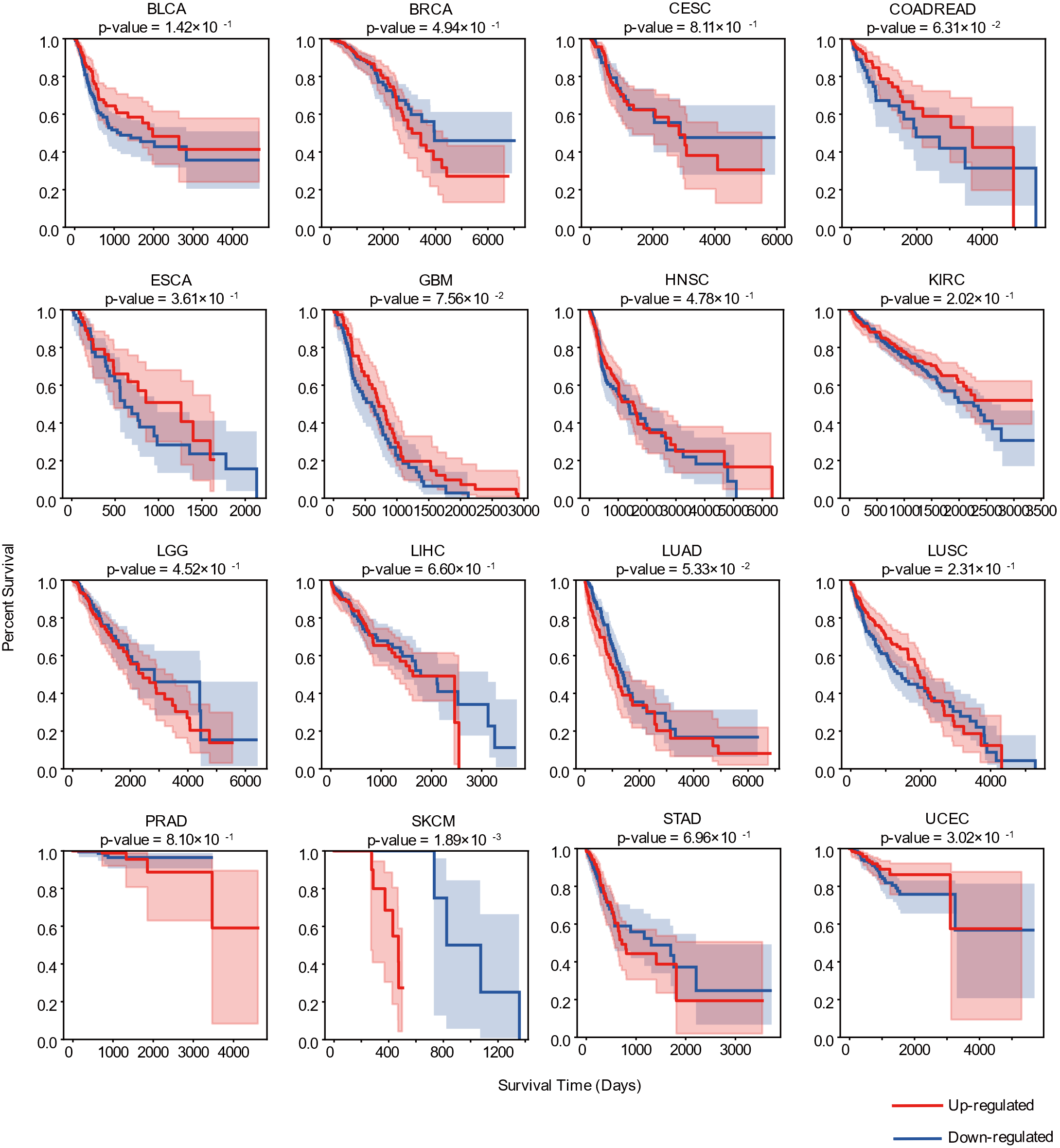

**Figure S14. Survival plot of patient subgroups in 16 cancer types stratified by driver genes of each cancer type.** Red lines represent patient groups with upregulated driver gene expression. Blue lines represent patient groups with downregulated driver gene expression. Significance of survival difference between patient groups is shown as the p-value of log-rank test.

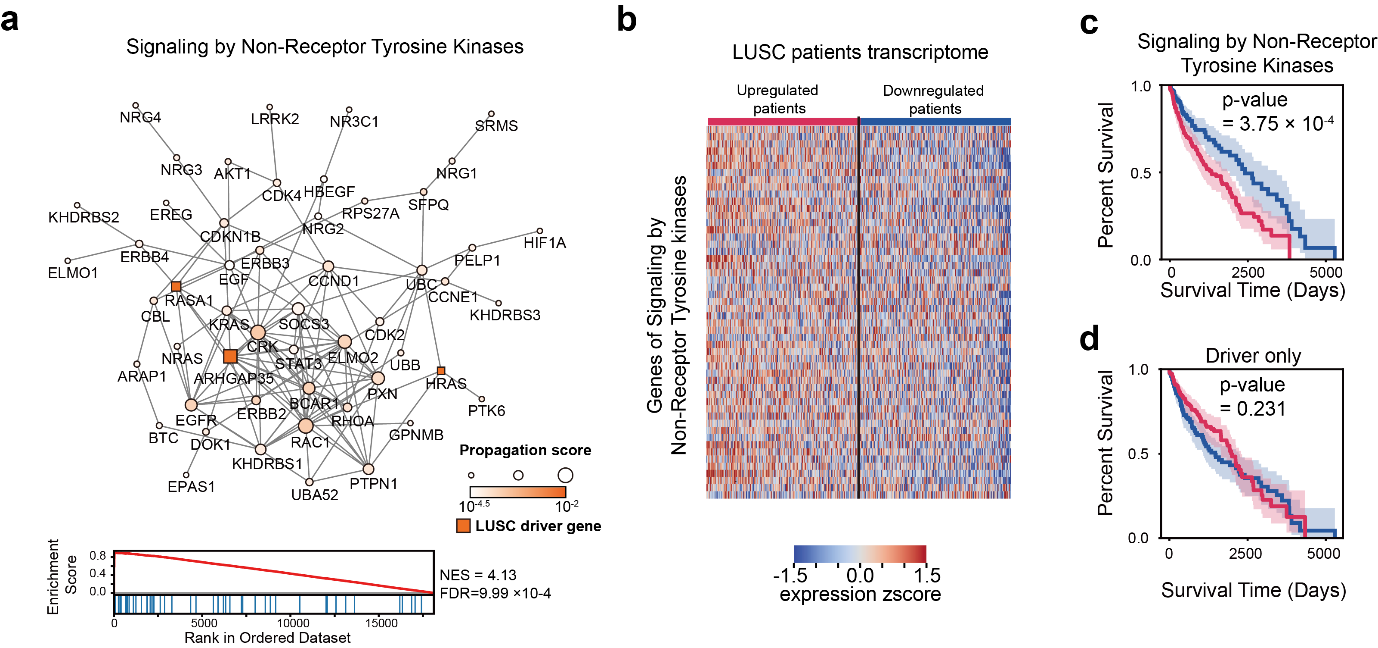

**Figure S15. Patient stratification using the co-essentiality network module (Signaling by Non-Receptor Tyrosine Kinases pathway) in LUSC. (a)** Subnetwork of the co-essentiality network for signaling by nonreceptor tyrosine kinase pathway (up). Enrichment plot of the co-essentiality network resulting from GSEA on Signaling by Non-Receptor tyrosine Kinase pathway (bottom). **(b)** Heatmap of LUSC gene expression in Signaling by Non-Receptor Tyrosine Kinases pathway. **(c)** Survival plot of LUSC patient subgroups stratified by gene expression of signaling by the nonreceptor tyrosine kinase pathway. **(d)** Survival plot of LUSC patient subgroups stratified by LUSC driver genes. Red and blue lines represent the upregulated and downregulated patient groups, respectively.

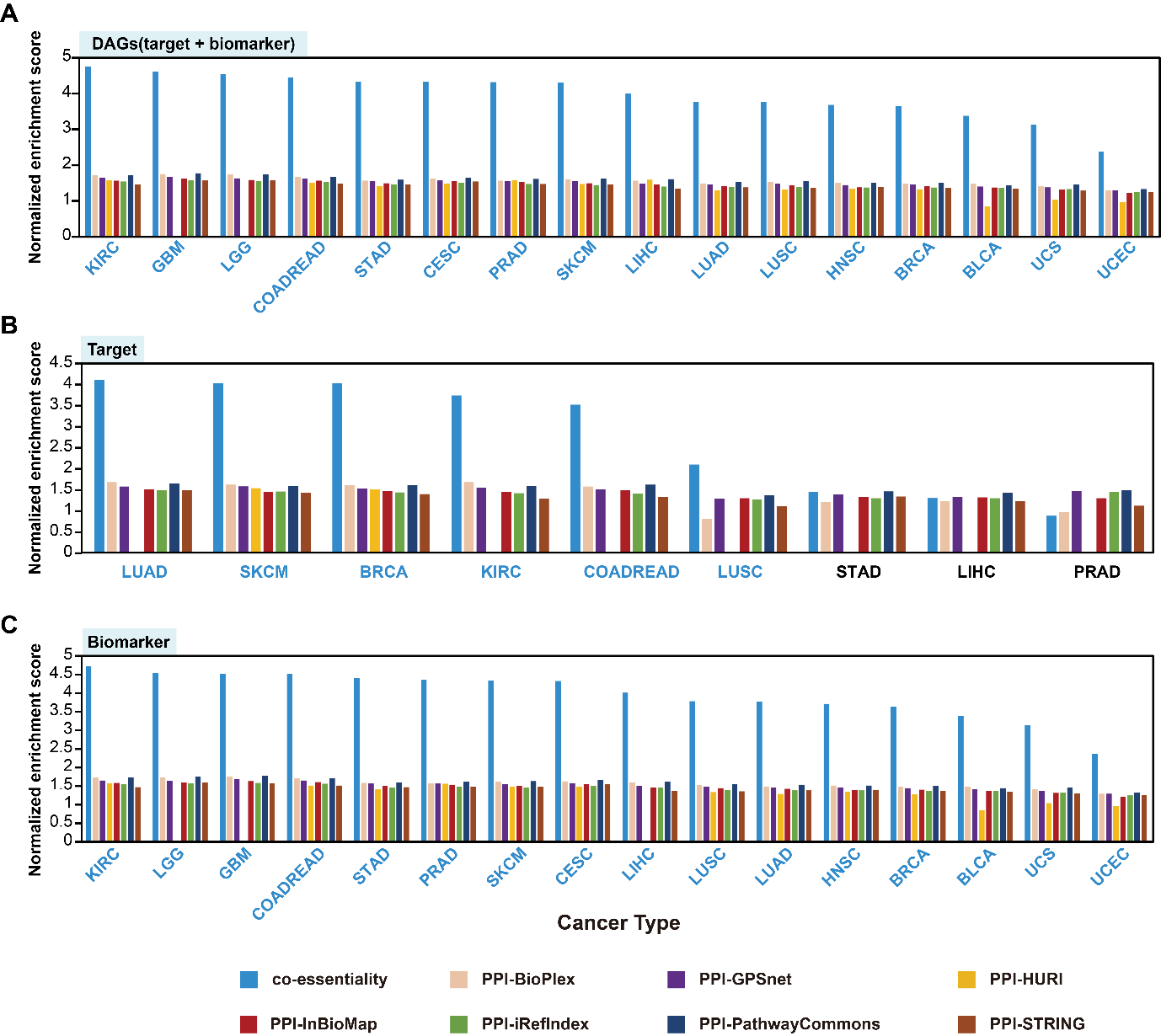

**Figure S16. Performance of the co-essentiality network for approved anticancer drug-associated gene(DAG) prioritization compared with that of seven PPI networks.** (**A**) The performance for anticancer DAG prioritization was measured using the normalized enrichment score (NES) for eight networks: co-essentiality, BioPlex, GPSnet, HURI, Inbiomap, iRefIndex, Pathway Commons, and STRING. **(B**) The prioritization performance for targets of approved anticancer drug was measured using the NES. **(C**) The prioritization performance for biomarkers of approved anticancer drug was measured using the NES. For blue-colored cancer types, the co-essentiality network showed the highest NES among the eight networks.

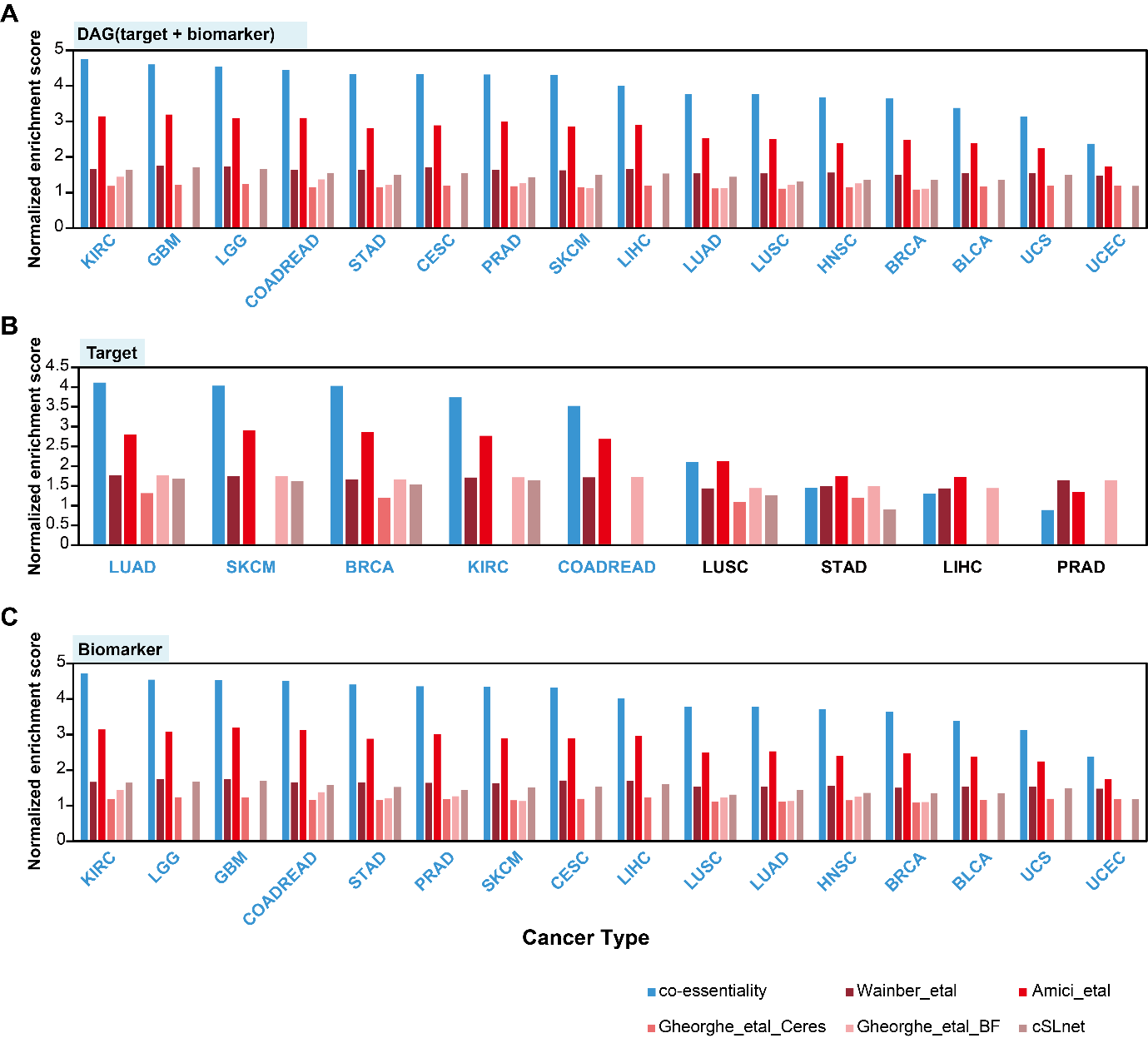

**Figure S17. Performance of the co-essentiality network for approved anticancer drug-associated gene(DAG) prioritization compared with that of other co-essentiality networks and a genetic interaction network.** (A) The performance for anticancer DAG prioritization was measured using the normalized enrichment score (NES) for other co-essentiality networks (Wainberg_etal, Amici_etal, Gheorghe_etal_Ceres, and Gheorghe_etal_BF), and the genetic interaction network(cSLnet). (B) The prioritization performance for targets of approved anticancer drug was measured using the NES. (C) The prioritization performance for biomarkers of approved anticancer drug was measured using the NES. For blue-colored cancer types, the co-essentiality network showed the highest NES among the six networks.

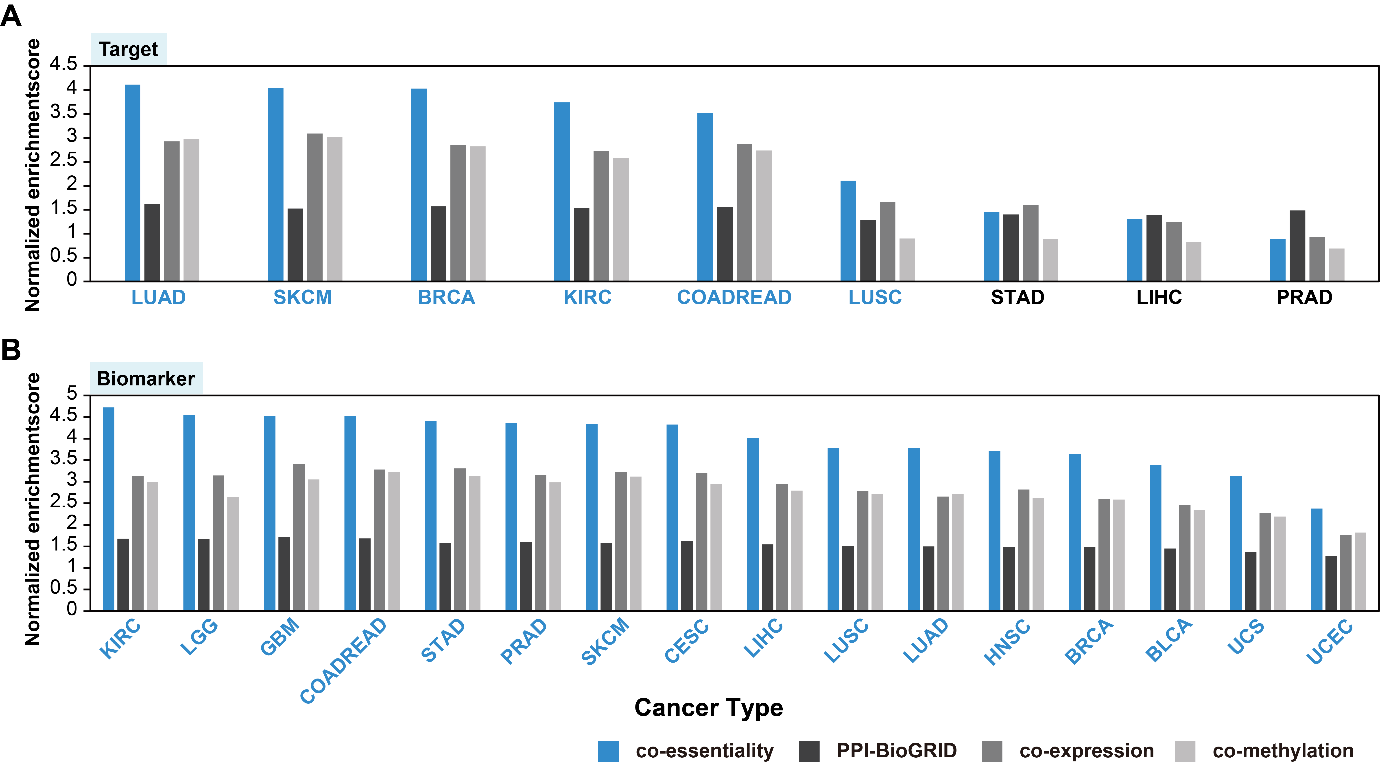

**Figure S18. Performance of the co-essentiality network for targets and biomarkers of approved anticancer drugs prioritization compared with that of the four networks.** (A) The prioritization performance for targets of approved anticancer drug was measured using the normalized enrichment score (NES) for four networks: co-essentiality, BioGRID, co-expression, co-methylation. (B) The prioritization performance for biomarkers of approved anticancer drug was measured using the NES. For blue-colored cancer types, the co-essentiality network showed the highest NES among the four networks.

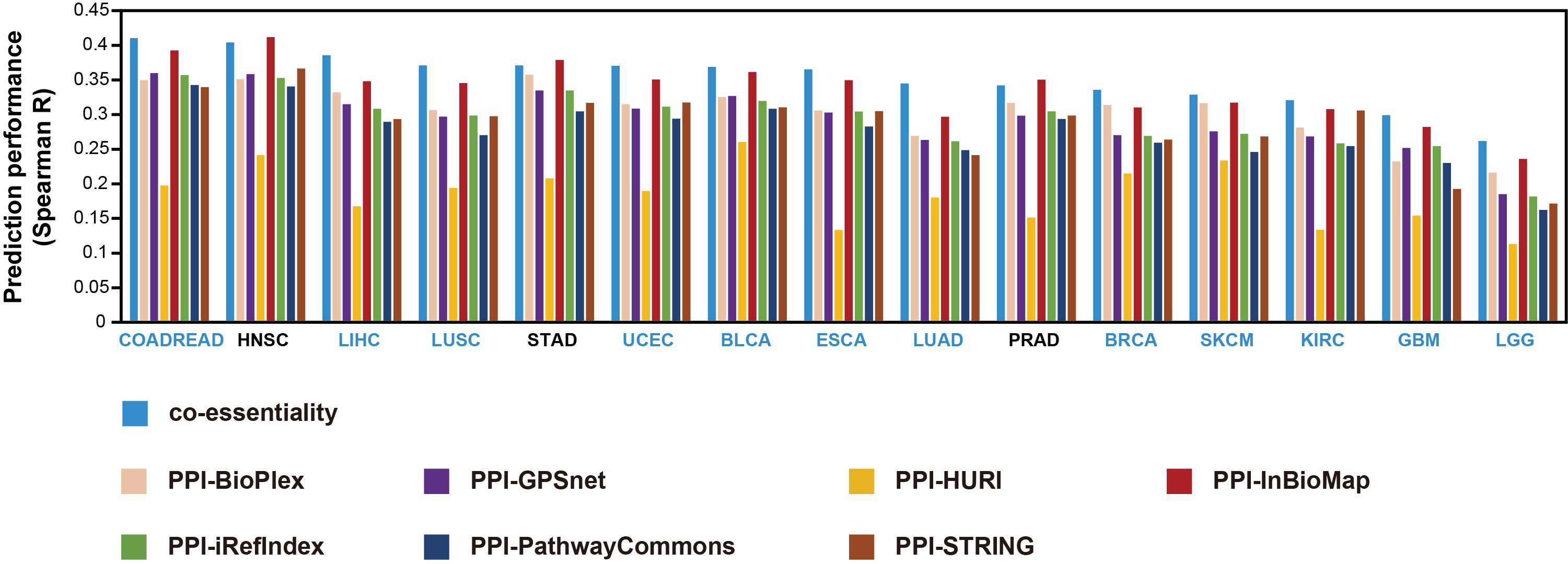

**Figure S19. Performance of the co-essentiality network for drug response prediction compared with that of seven different PPI networks.** The performance of drug response prediction was measured using the Spearman correlation coefficient (Spearman R) for eight networks: co-essentiality, BioPlex, GPSnet, HURI, Inbiomap, iRefIndex, Pathway Commons, and STRING. For blue-colored cancer types, the co-essentiality network showed the best performance among the eight networks.

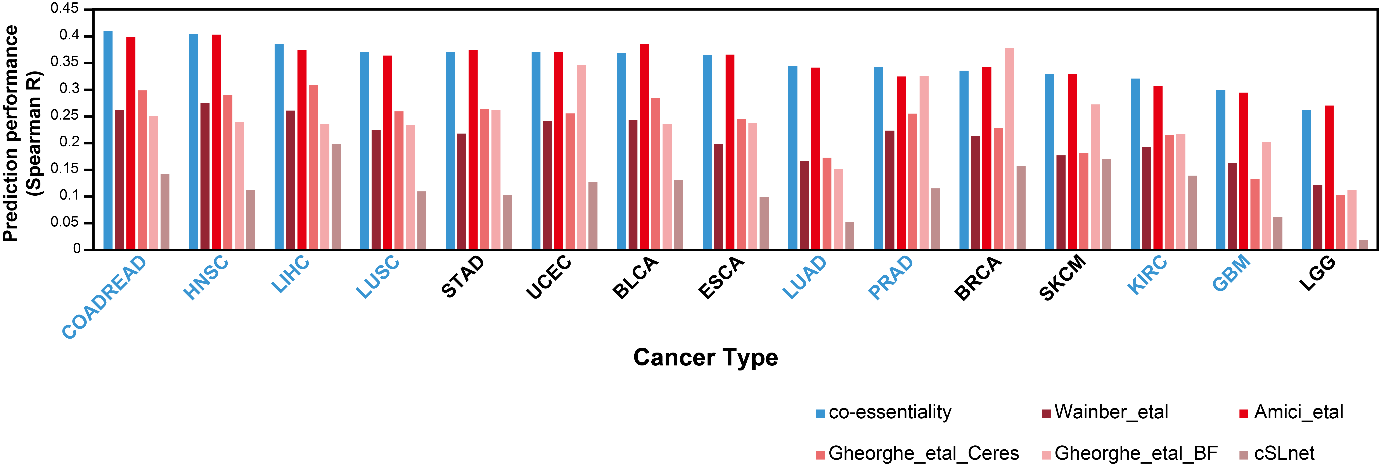

**Figure S20. Performance of the co-essentiality network for drug response prediction compared with that of another co-essentiality network and genetic interaction network.** The performance of drug response prediction was measured using the Spearman correlation coefficient (Spearman R) for other co-essentiality networks (Wainberg_etal, Amici_etal, Gheorghe_etal_Ceres, and Gheorghe_etal_BF), and the genetic interaction network (cSLnet). For blue-colored cancer types, the co-essentiality network showed the best performance among the six networks.

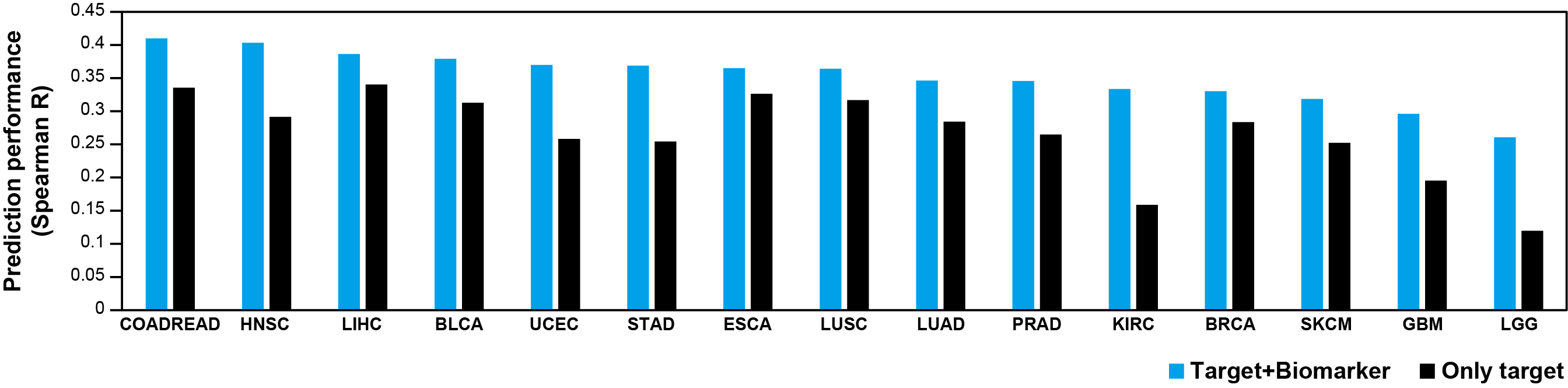

**Figure S21. Comparing the performance of drug response prediction using both target and biomarker information versus using only target information.** Blue colored bars are prediction performance of drug response using the TC score based on DAGs including both targets and biomarkers. Black colored bars are prediction performance of drug response using the TC score based on only target information.

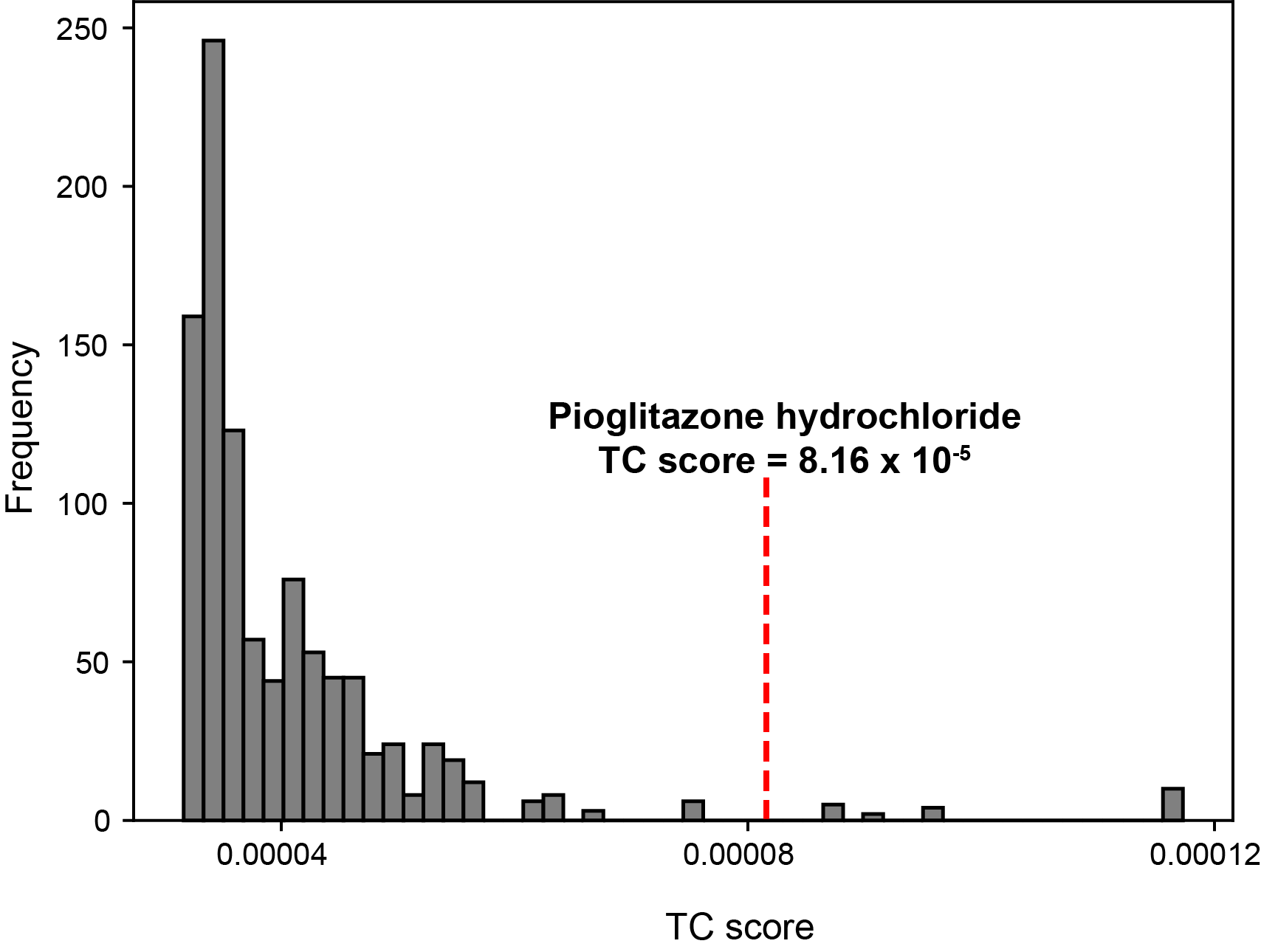

**Figure S22. Empirical distribution of the TC score of the degree controlled random nodes for drug-associated genes(DAGs) of Pioglitazone hydrochloride.** The x-axis shows the value of the TC score, while the y-axis denotes the fraction of the number of random DAGs against 100 permutations. The red dot line is the observed TC score of Pioglitazone hydrochloride.

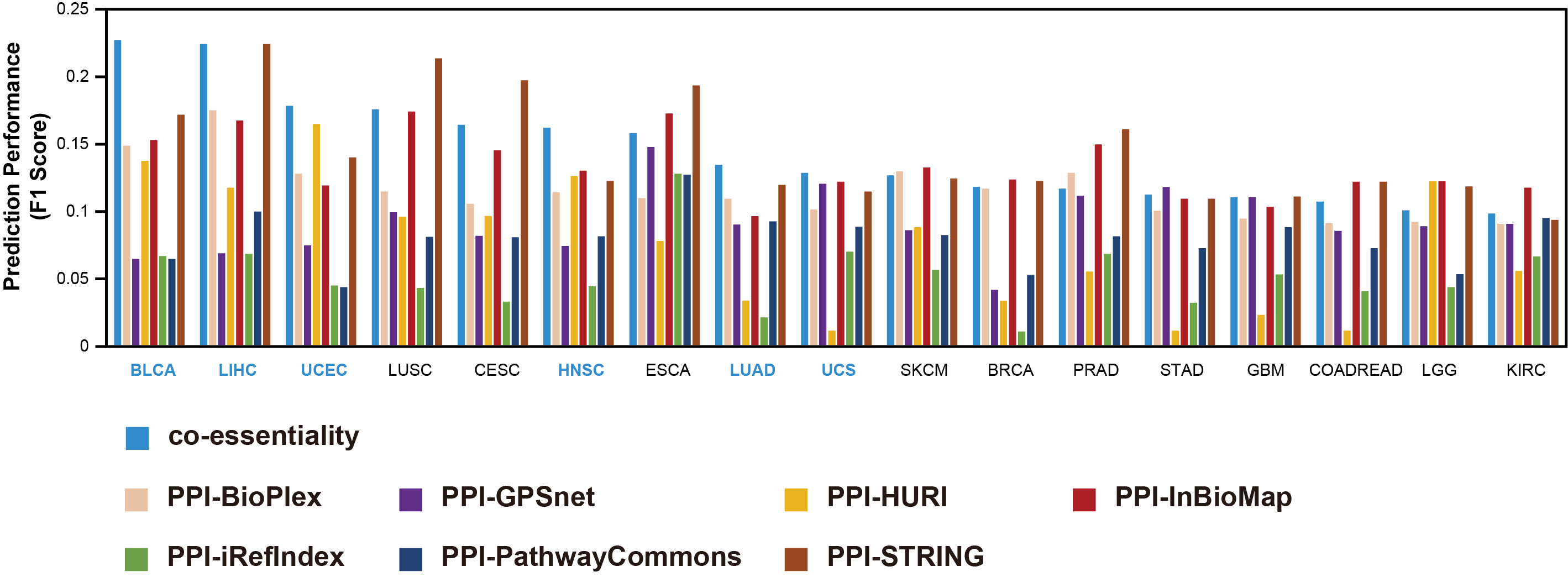

**Figure S23. Performance of the co-essentiality network for repurposing of non-cancer approved drugs compared with that of seven different PPI networks.** The performance for repurposing of non-cancer-approved drugs was measured using the F1 score for eight networks: co-essentiality, BioPlex, GPSnet, HURI, Inbiomap, iRefIndex, Pathway Commons, and STRING. For blue-colored cancer types, the co-essentiality network showed the highest F1 score among the eight networks.

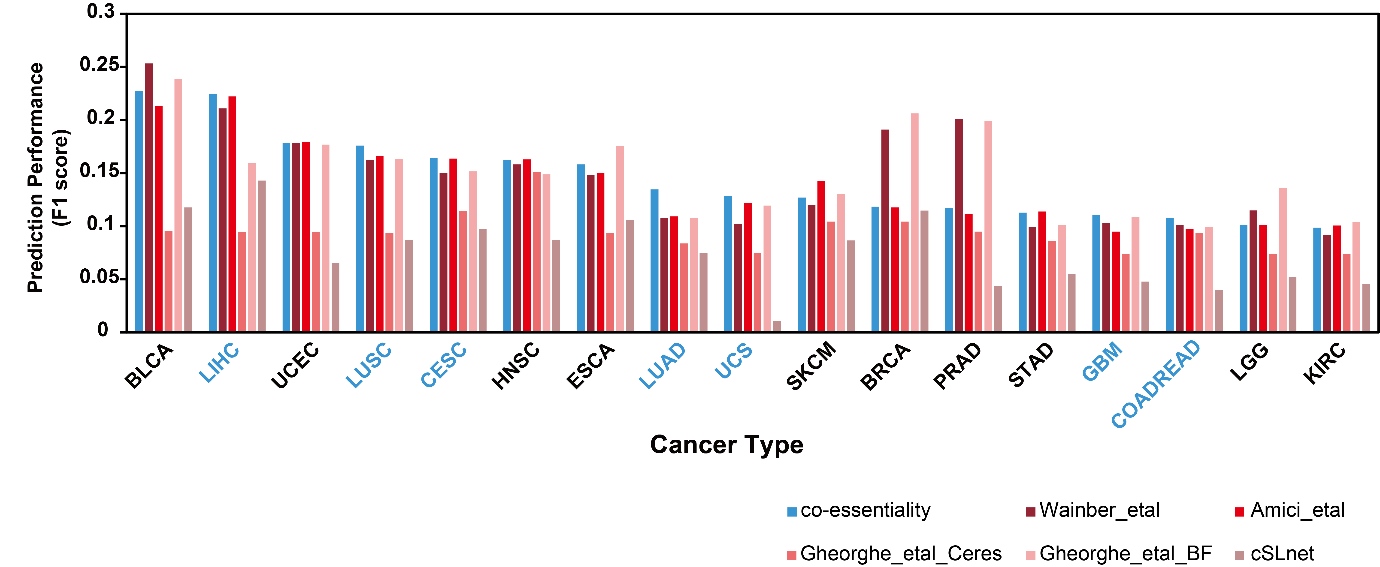

**Figure S24. Performance of the co-essentiality network for repurposing of non-cancer approved drugs compared with that of other co-essentiality networks and a genetic interaction network.** The performance for repurposing of non-cancer approved drugs was measured using the F1 score for other co-essentiality network (Wainberg_etal, Amici_etal, Gheorghe_etal_Ceres, and Gheorghe_etal_BF), and a genetic interaction network (cSLnet). For blue-colored cancer types, the co-essentiality network showed the highest F1 score among the three networks.

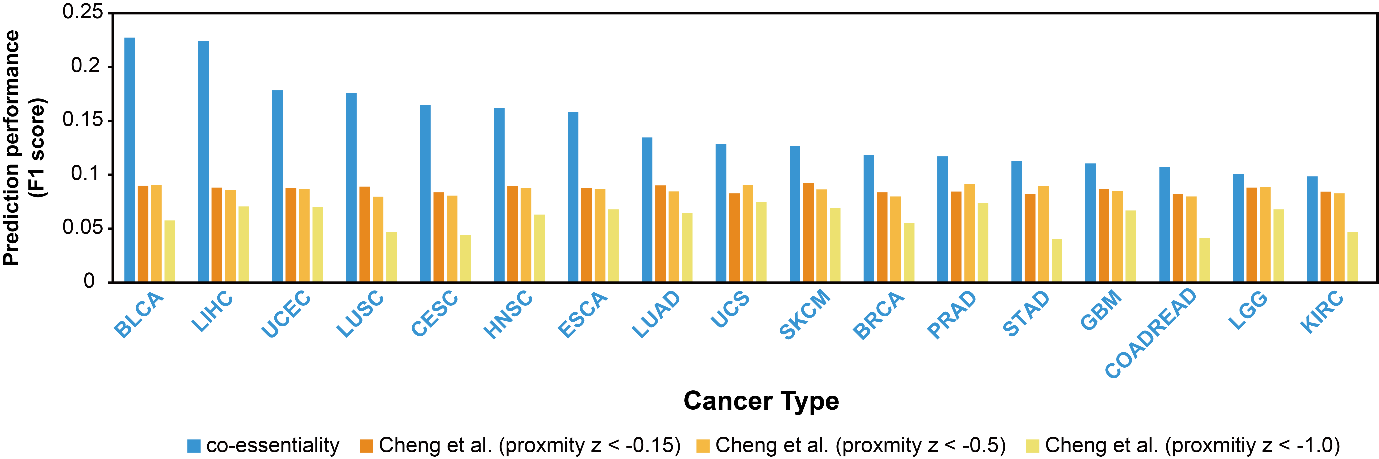

**Figure S25. Comparison of in silico drug repurposing performance between co-essentiality network method and Cheng et al. method.** Drug repurposing performance was evaluated using clinical trial records as validation. Prediction performance was measured using the F1 score. Yellow spectrum bars indicate the performance of the Cheng et al. method at three different network proximity thresholds for identifying repurposed drugs. For cancer types shown in blue, the co-essentiality network demonstrated superior performance compared to all three alternative approaches.

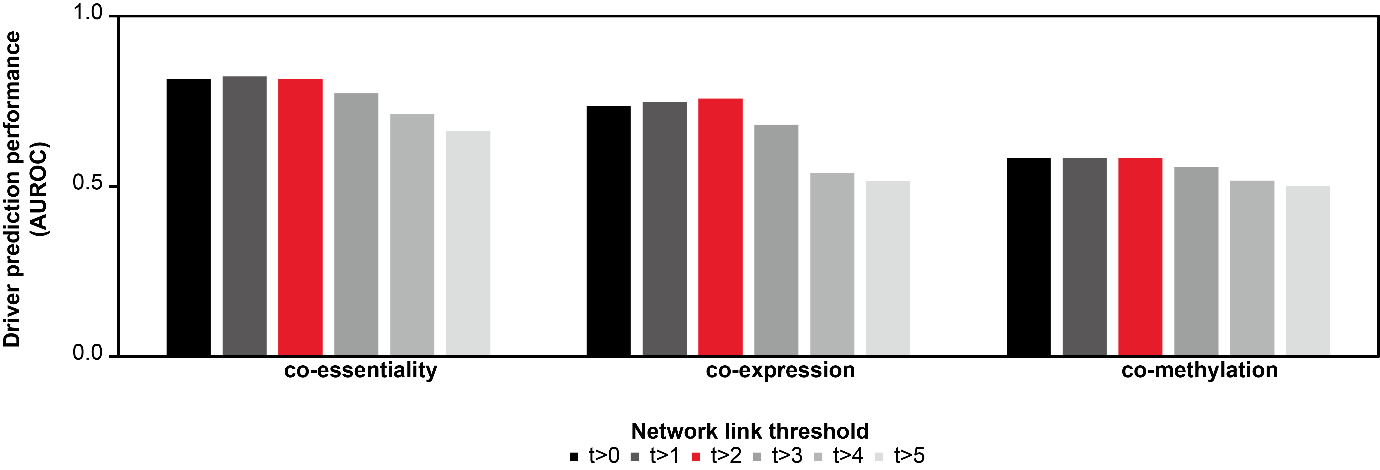

**Figure S26. Performance comparison of cancer driver gene identification across three correlation-based networks (co-essentiality, co-expression, and co-methylation) using different network link thresholds ranging from 0.0 to 5.0.** Red bars indicate results at the selected threshold t > 2.0 used in this study.

**Table S2.** Information of the 16 networks used in this study.

| **Network** | **Node**  **number** | **Link**  **number** | **Essential gene**  **number** | **Essential gene**  **fraction** | **Rank of**  **fraction** |
| --- | --- | --- | --- | --- | --- |
| co-essentiality | 18119 | 8105180 | 2070 | 11.4% | 9 |
| PPI-BioGRID | 18708 | 434527 | 2011 | 10.7% | 13 |
| co-expression | 19120 | 12403225 | 2120 | 11.1% | 11 |
| co-methylation | 16333 | 10193089 | 1732 | 10.6% | 14 |
| PPI-BioPlex | 12848 | 86968 | 1700 | 13.2% | 3 |
| PPI-GPSnet | 15124 | 167854 | 1948 | 12.9% | 5 |
| PPI-HURI | 8124 | 51816 | 1009 | 12.4% | 6 |
| PPI-Inbiomap | 17421 | 608161 | 2014 | 11.6% | 7 |
| PPI-iRefIndex | 14955 | 152147 | 1930 | 12.9% | 4 |
| PPI-PathwayCommons | 17646 | 88329 | 2035 | 11.5% | 8 |
| PPI-STRING | 18119 | 770875 | 2070 | 11.4% | 9 |
| Wainberg_etal | 6351 | 14980 | 1191 | 18.8% | 1 |
| Amici_etal | 19082 | 1040194 | 2089 | 10.9% | 12 |
| Gheorghe_etal_Ceres | 12910 | 359564 | 1847 | 14.3% | 2 |
| Gheorghe_etal_BF | 2025 | 6244 | 210 | 10.4% | 15 |
| cSLnet | 8368 | 21450 | 741 | 8.9% | 16 |

**Table S4.** FDA-approved drug-associated genes (DAGs) of SKCM among the top 50 genes with the highest propagation values in the co-essentiality network.

| **Rank #** | **Gene** | **Driver** | **Direct Target** | **Biomarker** |
| --- | --- | --- | --- | --- |
| 1 | TP53 | O |  | VEMURAFENIB, PACLITAXEL, DABRAFENIB, VORINOSTAT, 5-FLUOROURACIL, TRAMETINIB |
| 2 | BRAF | O | DABRAFENIB, VEMURAFENIB | PACLITAXEL, VORINOSTAT, 5-FLUOROURACIL, TRAMETINIB, COBIMETINIB, DABRAFENIB MESYLATE |
| 3 | RAC1 | O |  | DABRAFENIB, VEMURAFENIB |
| 5 | CTNNB1 | O |  | TRAMETINIB |
| 7 | KRAS | O |  | VEMURAFENIB, PACLITAXEL, DABRAFENIB, 5-FLUOROURACIL, TRAMETINIB, COBIMETINIB |
| 8 | RB1 | O |  | VORINOSTAT, TRAMETINIB |
| 10 | PTEN | O |  | PACLITAXEL, VORINOSTAT, VEMURAFENIB |
| 11 | NF1 | O |  | DABRAFENIB, COBIMETINIB, VEMURAFENIB, TRAMETINIB |
| 12 | MAP2K1 | O | COBIMETINIB, TRAMETINIB | DABRAFENIB MESYLATE, DABRAFENIB, VEMURAFENIB, COBIMETINIB FUMARATE, 1187431-43-1 |
| 14 | NRAS | O |  | DABRAFENIB, COBIMETINIB, VEMURAFENIB, 5-FLUOROURACIL, TRAMETINIB |
| 18 | HRAS | O |  | TRAMETINIB |
| 20 | KIT | O |  | TRAMETINIB |
| 22 | GNA11 | O |  | VORINOSTAT, TRAMETINIB |
| 23 | CDKN2A | O |  | PACLITAXEL, TRAMETINIB |
| 27 | CDKN1A | X |  | PACLITAXEL, 5-FLUOROURACIL |
| 30 | ATM | X |  | PACLITAXEL, TRAMETINIB |
| 32 | CRKL | X |  | DABRAFENIB, VEMURAFENIB |
| 34 | SOX10 | X |  | VEMURAFENIB |
| 40 | SOX9 | X |  | 5-FLUOROURACIL |
| 47 | RAF1 | X | DABRAFENIB | VEMURAFENIB |

**Table S6.** Seven repurposing candidates in LUAD predicted by the co-essentiality network but not by other networks.

| **Drug** | **TC score** | **Adjusted P** | **clinical trial records in cancer** |
| --- | --- | --- | --- |
| PIOGLITAZONE HYDROCHLORIDE | $8.16\times{10}^{-5}$ | $3.79\times{10}^{-3}$ | O |
| ROSIGLITAZONE MALEATE | $8.16\times{10}^{-5}$ | $3.79\times{10}^{-3}$ | X |
| ATOVAQUONE | $7.81\times{10}^{-5}$ | $2.57\times{10}^{-3}$ | O |
| INSULIN GLARGINE | $7.68\times{10}^{-5}$ | $1.23\times{10}^{-4}$ | X |
| TERIFLUNOMIDE | $7.42\times{10}^{-5}$ | $1.09\times{10}^{-2}$ | X |
| EFLORNITHINE | $6.73\times{10}^{-5}$ | $1.77\times{10}^{-2}$ | O |
| TELMISARTAN | $6.67\times{10}^{-5}$ | $3.95\times{10}^{-2}$ | X |

**Table S7.** AUROC of driver gene identification of three correlation-based networks according to six threshold values(t).

| **Threshold t** | **co-essentiality** | **co-expression** | **co-methylation** |
| --- | --- | --- | --- |
| 0 | 0.815 | 0.737 | 0.583 |
| 1 | 0.823 | 0.747 | 0.583 |
| 2 | 0.814 | 0.758 | 0.583 |
| 3 | 0.773 | 0.681 | 0.556 |
| 4 | 0.711 | 0.539 | 0.516 |
| 5 | 0.662 | 0.516 | 0.502 |

**Table S8.** Comparison of AUROC for driver gene identification using the correlation-based and mutual information-based co-expression networks across six threshold values (t).

| **Threshold t** | Pearson correlation coefficient  (PCC) | Maximal information coefficient  (MIC) |
| --- | --- | --- |
| 0 | 0.737 | 0.611 |
| 1 | 0.747 | 0.648 |
| 2 | 0.758 | 0.721 |
| 3 | 0.681 | 0.698 |
| 4 | 0.539 | 0.635 |
| 5 | 0.516 | 0.580 |

**Table S1.** Relative modularity of 186 KEGG pathways on four networks and the cancer-related pathway (CRP) list (Separated Excel file)

**Table S3.** List of FDA approved anticancer drugs and their corresponding DAGs (Separated Excel file)

**Table S5.** List of drug repurposing candidates predicted by the co-essentiality network (Separated Excel file)

**Table S9.** ‘Tier 1’ driver genes in the cancer gene census(CGC) and matched TCGA cancer types (Separated Excel file)

**Table S10.** Drug-gene association and drug information (Separated Excel file)
